# The dorsolateral striatum encodes a temporal basis for the organization of behavior

**DOI:** 10.1101/2023.11.13.566826

**Authors:** Filipe S. Rodrigues, Tiago Monteiro, Asma Motiwala, Joseph J. Paton

## Abstract

To behave adaptively, the brain must register temporal structure in the environment and use it to organize behavior. The dorsolateral striatum (DLS) integrates sensorimotor input, and is necessary for accurate timing and structuring behavior in general. However, if DLS provides the basis for mapping temporal features in the environment to behavior, its activity should predict variation in that mapping. A reanalysis of DLS population activity in rats comparing the duration of two sequentially presented vibratory stimuli revealed a striking correspondence between neural activity and behavior. Varying vibration intensity of the second stimulus induced systematic biases in temporal judgments, and corresponding biases in multiple features of DLS activity during stimulus presentation, including population coding of time. In contrast, the same intensity manipulations applied to the first stimulus affected neither behavior nor neural activity. Furthermore, neuronal response profiles were best described as a continuum, arguing against hypotheses where categories of responses, e.g., ramping activity, selectively underpin temporal processing. These data represent important additional evidence that striatal population dynamics support the organization of behavior by mapping temporal information to action.

## Introduction

Evidence for the involvement of striatum in interval timing comes from a variety of sources, including disease states (Malapani et al., 1998; Rowe et al., 2010), lesions (Meck, 2006; Wolff et al., 2022), neuroimaging (Coull et al., 2004; Rao et al., 2001), neurophysiology (Gouvêa et al., 2015; Jin et al., 2009; Kim et al., 2018; Matell et al., 2003; Mello et al., 2015; Monteiro et al., 2023), pharmacology (Gouvêa et al., 2015; Maricq & Church, 1983; Mello et al., 2015), optogenetics (Cruz et al., 2022) and temperature manipulations (Monteiro et al., 2023). A recent study challenged this body of work by concluding that though time information is inherent to neural activity in the DLS, it is otherwise decoupled from animals’ estimates of elapsed time, as inferred from their duration judgments (Toso, Reinartz, et al., 2021). The data used to support this claim was recorded during two variants of a delayed comparison task. Briefly, rats were presented with two sequential vibratory stimuli that varied in both duration and intensity, and trained to either compare the duration and ignore the intensity (*delayed duration comparison*), or compare the intensity and ignore the duration (*delayed intensity comparison*) of the two stimuli. The authors’ central conclusion that striatal activity was decoupled from temporal judgment was based on four purported features of DLS activity:

1. though rats’ judgments about the duration of vibration stimuli were reliably biased by their intensity, activity in the DLS was not similarly modulated;
2. the DLS did not encode stimulus duration differently on correct and incorrect trials;
3. elapsed time was decodable from DLS activity in all epochs of the delayed duration comparison task, and not just during stimulus periods;
4. in rats trained in the delayed intensity comparison task variant, stimulus duration was encoded by DLS even though it was experimentally irrelevant.

Here, we focus on the delayed duration comparison task to demonstrate that 1) and 2) are false – DLS activity was unambiguously modulated by stimulus intensity and predictive of animals’ duration judgments. This was readily apparent in the population mean activity when conditioned on intensity or choice and corroborated by multiple quantitative analyzes, including state space visualization, the construction of neurometric curves, probabilistic population decoding of elapsed time and finally a generalized linear model (GLM) that seeks to partition the relative contribution of different task variables to explainable variance in neural activity. We also argue that 3) and 4) logically do not constitute support for the claim that striatal activity is not used to guide judgment of duration. By analogy, would the response of a primary visual cortex (V1) neuron to a visual cue incidentally presented within its receptive field, but without behavioral importance for some task at hand lead one to conclude that V1 does not support visual behavior? Just as we do not go intermittently blind in the absence of an explicitly visual task at hand, we would not expect timing mechanisms to shut off whenever we are not tasked with judging the passage of time.

In addition, we detail the several data analysis problems that lead us to question the central conclusions of the original study and in its erratum. Upon addressing these issues, we find the familiar correlation between speed of striatal dynamics and judgments about duration, as well as a striking agreement between the degree to which different task variables impacted choices and were encoded by DLS neurons.

## Results

### Rats did not discriminate vibration durations as idealized and reported

Toso et al. performed chronic extracellular recordings from the left DLS of three rats trained in a delayed duration comparison task (Fig. 1A, B). Briefly, rats were rewarded for correctly reporting which of two vibratory stimuli delivered to the right vibrissae was of a longer duration. Each trial consisted of a series of delays: a post-initiation delay of 700 ms; the first stimulus (S1) period; an inter stimulus interval (ISI) lasting 2 s; the second stimulus (S2) period; and a 500-ms post-stimulus delay (Fig. 1B). Animals were required to maintain their snout at the stimulus delivery port from trial initiation up until an auditory “go” cue was presented at the offset of the post-stimulus delay, at which point they were free to report their choice at one of the two lateral spouts (Fig. 1A). Importantly, in addition to the two vibrations varying along the task-relevant axis of duration, they varied along the task-irrelevant axis of intensity. Thus, individual stimulus pairs were defined on the basis of their duration (T1 and T2) and intensity (I1 and I2), and chosen such that the sets of all possible durations T = {112, 161, 232, 334, 481, 694, 1000} ms and intensities I = {34, 64, 117} mm s^−1^ were the same for both stimuli (Hernández et al., 1997).

**Figure 1:**
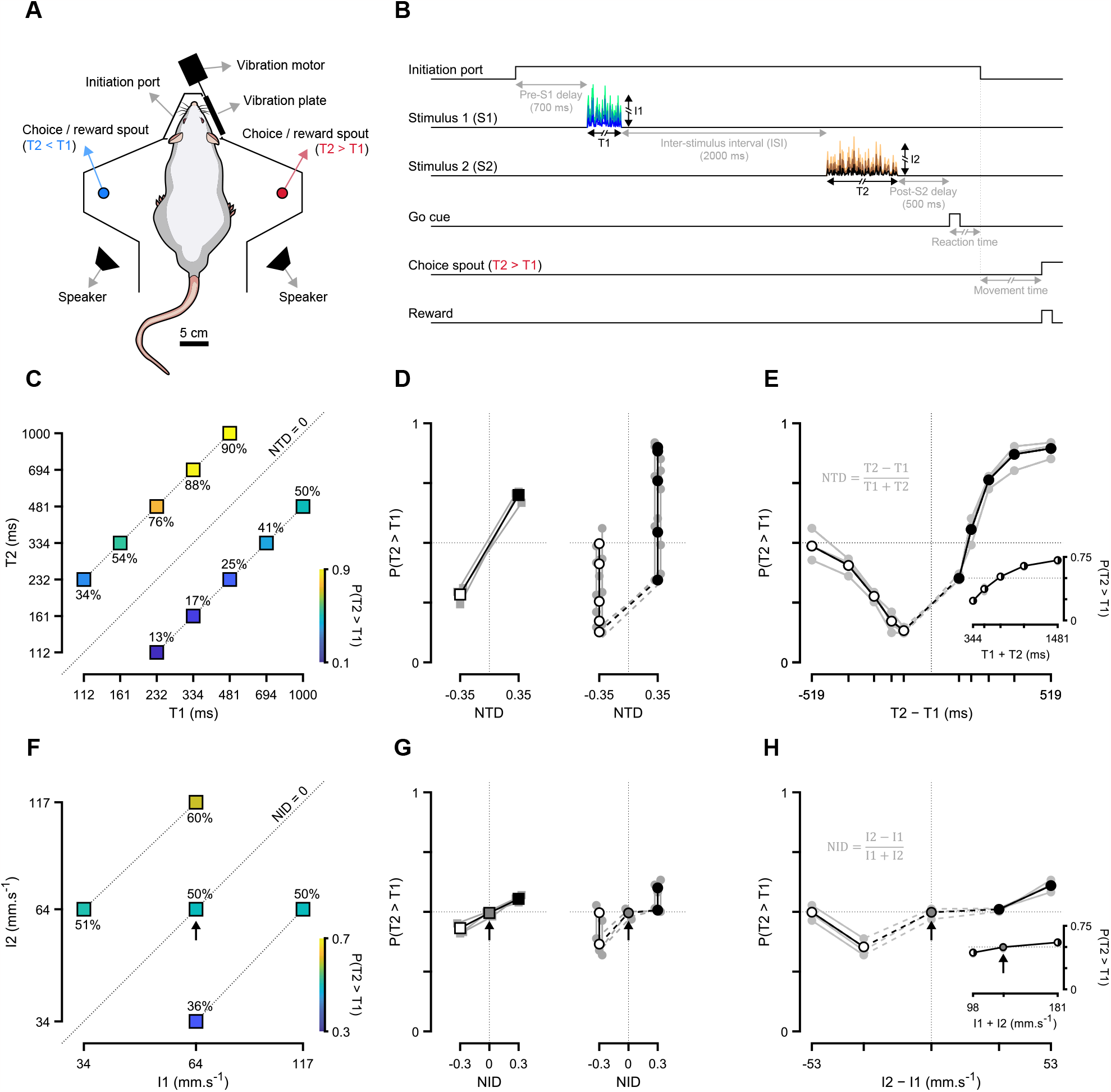
Rats did not discriminate vibration durations as idealized and reported. (**A**) Schematic of the behavioral apparatus with a Wistar rat depicted at the initiation/stimulus delivery port. The mapping of choice spout (left or right) to choice category (T2 > T1 or T2 < T1) was different from the depicted one for one of the three DLS-implanted rats. (**B**) Event diagram for an example correct trial in the delayed duration comparison task. (**C**) Proportion of T2 > T1 choices for all stimulus duration pairs pooled across the three DLS-implanted rats. Dotted lines above, below and at the identity line represent NTD levels of +0.35, −0.35 and 0, respectively. (**D**) Proportion of T2 > T1 choices as a function of NTD and grouped by NTD level (left) or stimulus pair (right). Gray markers and lines refer to individual animals, and black and white ones to cross-animal averages for each NTD level. Dashed lines on the right connect the two stimulus pairs with smaller absolute duration differences (|T2 − T1|). (**E**) Proportion of T2 > T1 choices conditioned on all possible values of the numerator (main axes) and denominator (inset) terms in the definition of NTD. (**F-H**) Same as (C-E), but for stimulus intensity and NID. The black arrow in (F-H) highlights the intensity pair that was undisclosed in both the original corrected versions of Toso, Reinartz, et al., 2021 (I1 = 64; I2 = 64 mm s^−1^).

Figure 1 shows all duration (Fig. 1C) and intensity pairs (Fig. 1F), as well as the corresponding performance pooled across the three DLS-implanted rats. Rather than annotating and/or using a diverging 3-hue color map to indicate *P*(T2 > T1), as done here (Fig. 1C, F) and in previous studies employing comparison tasks (Akrami et al., 2018; Fassihi et al., 2014; Romo et al., 1999), the authors denoted performance using only color in a sequential 2-hue color map. This makes it difficult to detect differences across stimulus pairs and, together with the authors’ undeclared omission of the most prevalent intensity pair in the data (I1 = 64; I2 = 64 mm s^−1^, Fig. S1), creates the impression that performance is well summarized as primarily driven by normalized time difference (NTD, Fig. 1D, left):

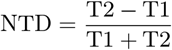

and biased by normalized intensity difference (NID, Fig. 1G, left):

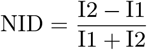

If animals’ choices were indeed independent of T1 and T2 within each NTD level, it follows that the probability of reporting T2 > T1 as a function of all possible values of T2 − T1 (the numerator in the definition of NTD) should produce a step-like function (Fig. S2). This was not the case in the data (Fig. 1D, E). The same rationale and prediction applies to NID and intensity difference (Fig. S2), and again the data did not conform (Fig. 1G, H). For both duration and intensity we show that *P*(T2 > T1) was a non-monotonic function of stimulus difference (Fig. 1E, H). Moreover, if animals’ decisions were truly governed by NTD and NID, then the denominators in their respective definitions, the aggregate stimulus duration (T1 + T2) and intensity (I1 + I2), should act only as normalization terms and as such not affect performance (Fig. S2). Instead, we found that rats’ propensity to judge T2 as larger than T1 increased monotonically as a function of both these quantities (insets in Fig. 1E, H).

Together, these observations reveal the inadequacy of NTD and NID as criteria for grouping trials when summarizing rats’ performance in this task and warrant further characterization of the behavior before searching for neural correlates of animals’ judgments.

### The second stimulus dominated the delayed duration comparison

We have shown that NTD and NID do not adequately capture performance in this task, and that the probability of rats reporting T2 as larger than T1 did not steadily increase as a function of the signed difference between these durations. Though this means that rats were not doing a one-to-one comparison of nominal stimulus durations, it is entirely consistent with them doing an asymmetric comparison due to what is commonly referred to as *contraction bias* or the *central tendency effect* – the tendency for perceptual judgments to gravitate towards the center of the distribution of previous stimuli. This behavioral regularity has been observed repeatedly across animal models and stimulus modalities (Akrami et al., 2018; Fassihi et al., 2014; Hollingworth, 1910; Jazayeri & Shadlen, 2010; Romo et al., 1999; Urai et al., 2017), and is proposed to arise from a Bayesian computation (Ashourian & Loewenstein, 2011; Xiang et al., 2021) wherein noisy representations of stimulus magnitudes are combined with prior information about the corresponding distribution of magnitudes (although see Boboeva et al., 2023 for an alternative non-Bayesian explanation). Indeed we find that rats’ performance was well accounted for by assuming that the effective magnitudes of T1 and I1 drifted towards the expected value of the respective sampling distributions during the inter-stimulus delay (Fig. 2A, B). The authors acknowledge that this phenomenon may have led to animals’ high error rate for the stimulus pair with the shortest possible T1 (T1 = 112; T2 = 232 ms, Fig. 2A), but in doing so underplay its ubiquity by suggesting that there is something unique about this particular stimulus duration pair.

**Figure 2:**
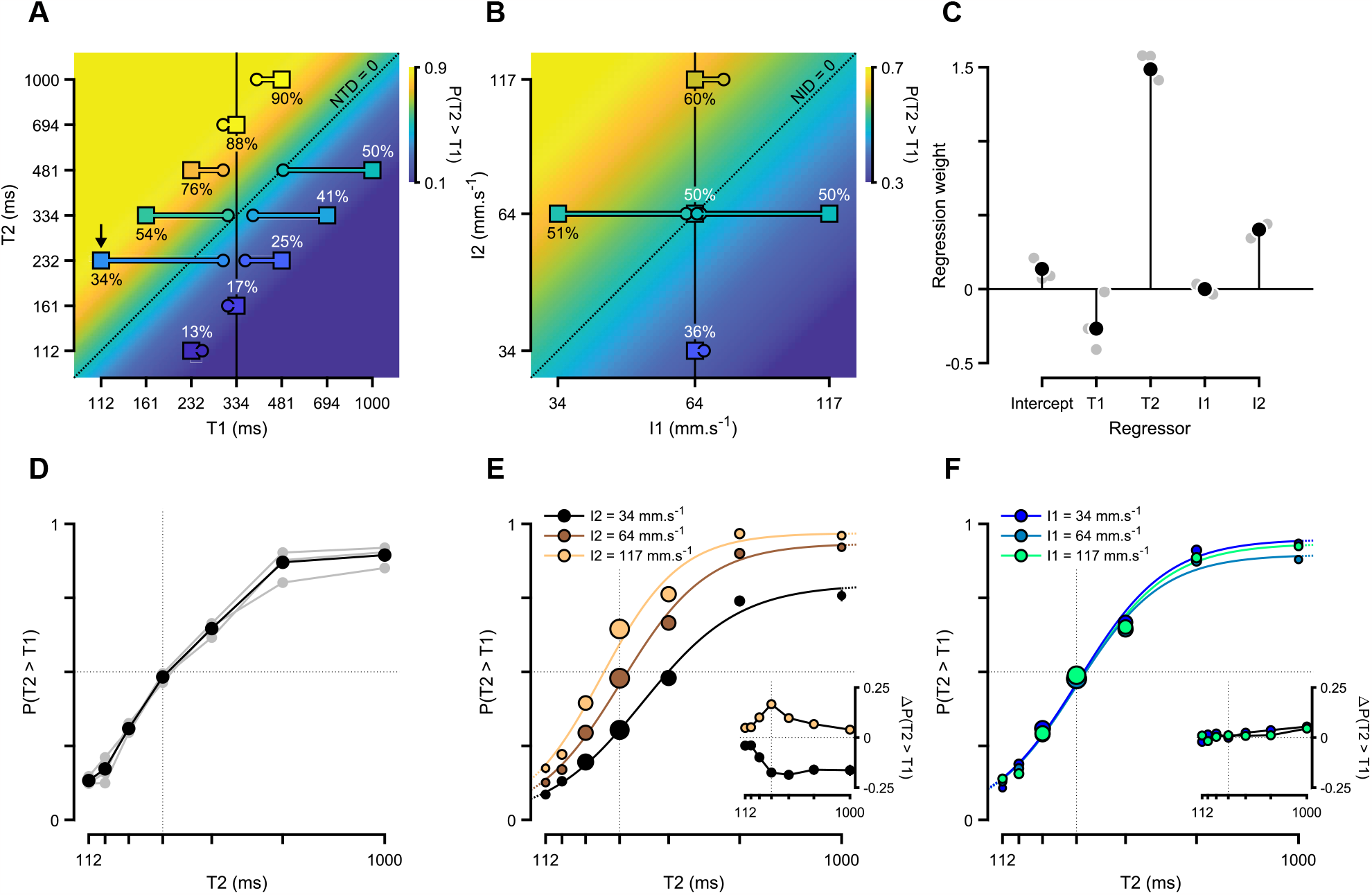
The second stimulus dominated the delayed comparison. (**A**) Depiction of how contraction bias may have affected estimates of T1 in the delayed duration comparison. Same data and analysis as in Fig. 1C overlaying a hypothetical performance rule where the probability of reporting T2 > T1 is a monotonic (sigmoidal) function of effective NTD. For each duration pair, a line was drawn between its T1 and T2 coordinates and the horizontally shifted point for which hypothetical performance was closest to rats’ performance on that pair. The vertical solid line depicts the expected value of the T1 distribution, and the black arrow indicates the one stimulus pair the authors singled-out as potentially affected by contraction bias. (**B**) Similar to (A), but for stimulus intensity. (**C**) Coefficients of a logistic regression model fit to choice data using all four stimulus features (T1, T2, I1 and I2) as regressors. Gray markers refer to fits for individual animals, and black ones to a fit pooling data across all three DLS-implanted animals. (**D**) Proportion of T2 > T1 choices as a function of T2. Gray markers refer to individual animals, and black ones data pooled across all three DLS-implanted animals. (**E**) Pooled discrimination performance as a function of T2 and conditioned on I2. Main axes: psychometric functions fit to I2-split choices. Marker size represents the relative number of trials per T2 value normalized within each I2 condition. Inset: differences between choice proportions in extreme-I2 (high: I2 = 117 mm s^−1^; low: I2 = 34 mm s^−1^) and intermediate-I2 trials (I2 = 64 mm s^−1^, mean ± propagated standard error of the mean (SEM)). (**F**) Same as (E), but conditioned on I1.

To quantify the asymmetry in the contribution of duration and intensity of the two stimuli onto animals’ judgments, we fit a logistic regression model using all four stimulus features (T1, T2, I1 and I2) as regressors and choice category (T2 > T1) as the response variable (Fig. 2C). Consistent with a strong contraction bias, T2 dominated the comparison, with an associated regression weight more than five times that of T1. Moreover, I1 had virtually no bearing on rats’ decisions, meaning that the intensity bias reported by the authors (Fig. 1G, left) was driven by I2 alone. Accordingly, plotting discrimination performance as a function of T2 produced monotonic functions (Fig. 2D), that systematically differed if conditioned on I2 (Fig. 2E) but not when conditioned on I1 (Fig. 2F). Lastly, fitting intensity-split choices with 4-parameter psychometric functions followed by model comparison confirmed the lack of detectable changes on parameters of psychophysical interest due to I1 and indicated that the effect of I2 could not be captured by any single psychometric parameter on its own (Fig. 2E, F, S3).

Beyond providing a more accurate summary of a rich behavior, these results inform subsequent analyzes of neural activity and constrain their interpretations. For example, lack of modulation of DLS neural activity by I1 takes on a different meaning after establishing that I1 did not meaningfully impact animals’ reports (Fig. 2C, F, S3) – the main behavioral covariate of internal estimates of elapsed time provided by the experimental paradigm. Likewise, the striking difference in the degree to which T1 and T2 predicted choices might warrant a similar asymmetry in emphasis on neural activity aligned to the first and second stimuli (Fig. 2C). Additionally, comparing neural activity recorded during nominally correct and incorrect trials is not readily interpretable due to the non-monotonic relationship between performance and experimental difficulty (Fig. 1E). Lastly, the fact that that extreme-I2 conditions did not simply produce laterally shifted, temporally or vertically rescaled versions of the intermediate-I2 psychometric curve (Fig. 2E, F, S3) argues against intensity affecting choices exclusively through any *one* property of the underlying decision variable (e.g., its temporal scale).

For these reasons, we focused most of the analyzes below on DLS activity recorded during the presentation of the second stimulus, refrained from using experimentally-defined correctness as a contrast when assessing if DLS responses covaried with duration judgments, and did not share in the authors’ assumption that temporal scaling of population dynamics is the only possible neural correlate of the observed I2-induced behavioral bias.

### Multiple features of DLS dynamics were systematically affected by the intensity of the second stimulus

Having established that rats’ behavior can be better described as categorization of T2 biased by I2 than discrimination of the two stimulus durations biased by their intensities, we next sought to characterize the impact of I2 on DLS activity at the single-cell and population levels during the second stimulus period. In their original publication, the authors reported that “*None of the neurons recorded* [in the DLS] *was found to encode I* [intensity] *by its firing rate*”, and this was later amended to “*2% of neurons encoded stimulus intensity during stimulus 2 presentation*” in their erratum. Viewing neural activity aligned to the onset of the second stimulus and conditioned on its intensity, both as population averages (Fig. 3A) and example neuronal responses (Fig. 3B), suggests a much higher-than-reported prevalence and strength of I2 modulation in the DLS. This is true in spite of cross-neuron averages being, in general, poor indicators of overall modulation, owing to their inability to distinguish between lack of modulation and symmetry in up- and down-modulation across the population. We will return later in the manuscript to this issue to explain why the authors reported so few neurons correlated with intensity.

**Figure 3:**
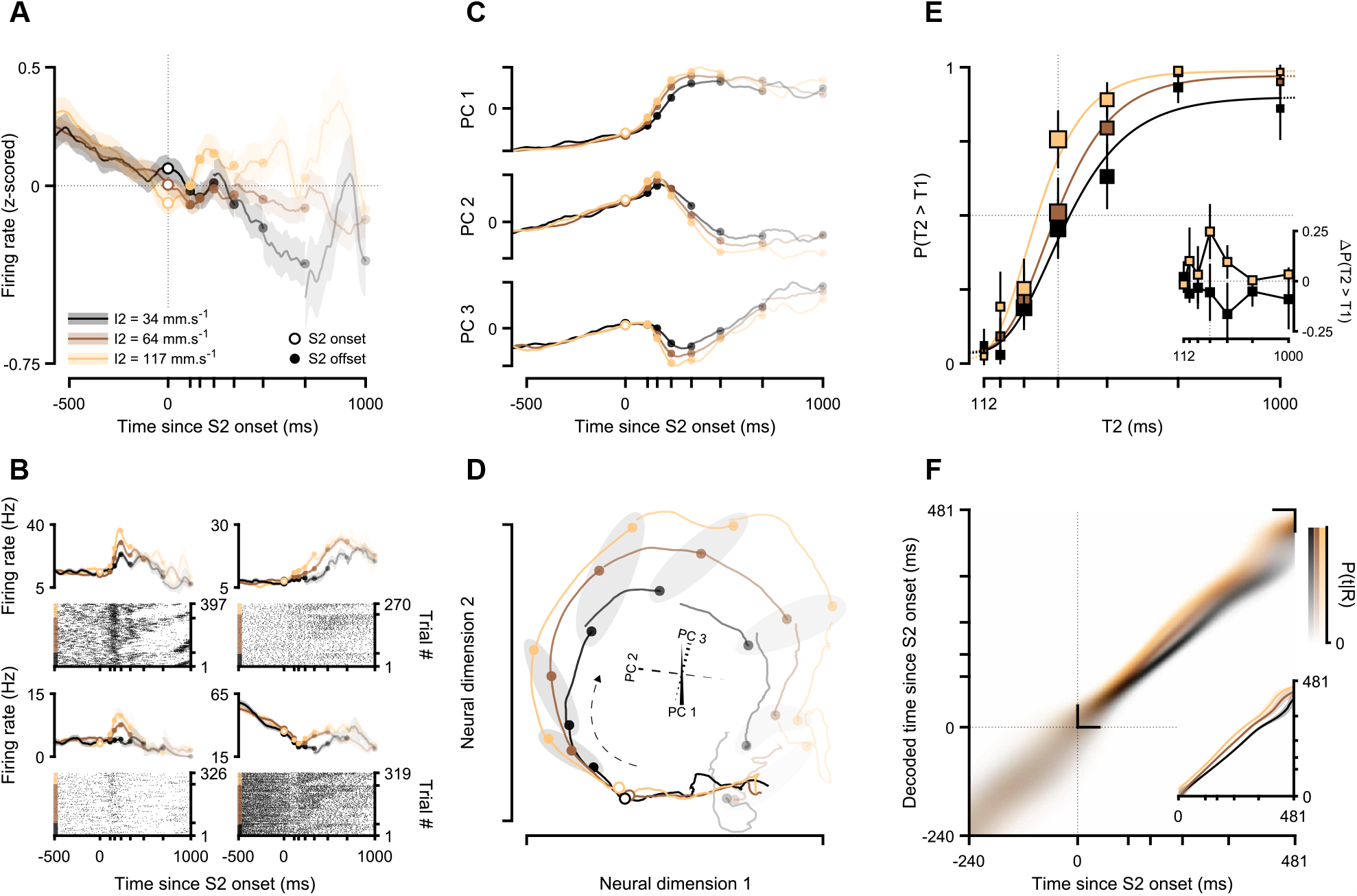
Multiple features of DLS neural activity were systematically affected by the intensity of the second stimulus. (**A**) Average normalized population activity conditioned on I2 and aligned to the onset of S2 (mean ± SEM across neurons). (**B**) Responses of four example DLS neurons during S2. For each neuron, spike density functions (mean ± SEM across trials) are conditioned on I2 and plotted above its spike raster, where trials are first sorted by I2, and then by T2 within each I2 level. (**C**) I2-split population activity projected onto each of the first three PCs computed using I2-agnostic population activity aligned to the onset of S2. (**D**) Population activity conditioned on I2 projected onto a 2D section of the subspace defined by the three PCs shown in (C). The 3-axis gizmo at the center depicts the projections of the axes defined by the first three PCs onto the chosen 2D section, the dashed black arrow indicates the direction of time, and the gray patches group population states recorded at each possible stimulus offset across I2. (**E**) Neurometric curves trained using S2-aligned population activity from middle-I2 trials conditioned on I2 (main axes), and differences between the proportion of T2 > T1 neural “reports” in high (I2 = 117 mm s^−1^) and low (I2 = 34 mm s^−1^) intensity trials compared to middle intensity (I1 = 64 mm s^−1^) trials (inset, mean ± propagated SEM). (**F**) Time estimates decoded from DLS activity around S2 presentation conditioned on I2. Main axes: mean posterior distributions across runs. Inset: *maximum a posteriori* (MAP) point estimates averaged across runs (median and IQR). The opacity of lines, markers and patches in (A-D) reflects the relative number of trials contributing to the underlying estimates (see Fig. 6D, bottom).

To further examine how I2 affected the time evolution of population activity of the DLS during the S2 period, we turned to dimensionality reduction using principal component analysis (PCA). Briefly, we computed principal components (PCs) using unconditioned S2-aligned average activity, and then projected average activity conditioned on I2 onto each of the first three PCs (Fig. 3C). This confirmed that a substantial amount of variance in DLS responses recorded during the presentation of the second stimulus is indeed explained by I2, and at least partially along the same intrinsic dimensions of neural space as T2-driven variance. Interestingly, I2 seemed to affect multiple features of DLS activity during this period, scaling it both in time and amplitude. This is already visible at the level of single-neuron responses (Fig. 3B) and PC projections (Fig. 3C), but is most evident when looking at neural trajectories in PC space (Fig. 3D), which seem to systematically vary with I2 in both speed and eccentricity. This dual effect of I2 on the temporal scale and overall gain of neural activity is reminiscent of its effect on multiple parameters of the psychometric function (Fig. 2E, S3), suggesting that the DLS may contribute to rats’ decisions in this task through dissociable features of its population dynamics.

Next, to characterize the contributions of intensity modulation of striatal activity onto the pattern of rats’ choices conditional on I2 and across T2, we performed cross-validated decoding of stimulus discrimination categories using neural population activity recorded at the end of S2 during middle-intensity trials. Specifically, we used half of all intermediate-I2 (I2 = 64 mm s^−1^) trials to train a linear classifier to discriminate the durations of the two stimuli (T1 and T2) based only on population spike counts recorded in the last 112 ms (duration of the shortest possible T2) before S2 offset. Neural “judgments” in all remaining trials (including all low- and high-intensity trials), were then simply the classes (T2 > T1 or T2 < T1) predicted by the classifier using those trials’ population state just prior to stimulus offset. The resulting neurometric curves split by I2 (Fig. 3E) provided a striking qualitative match to their psychometric counterparts (Fig. 2E), thus confirming that DLS population state nearing the offset of the second stimulus covaried with intensity-biased judgments for all possible stimulus durations.

Lastly, to characterize temporal scaling of population activity in DLS due to I2-driven modulation, we decoded elapsed time continuously from the population state during S2 presentation using a cross-validated naїve Bayes decoder. To do so, we first characterized the “typical” temporal profiles of striatal responses using a subset of all middle-intensity trials, and then applied a probabilistic decoding approach to estimate elapsed time based only on the observed state of the recorded population in remaining trials. Estimates of elapsed time derived from ongoing population activity systematically led ahead and lagged behind true time during high- and low-I2 trials, respectively (Fig. 3F). This decoding approach is something the authors tried as well, though with a different outcome and subsequent interpretation. The discrepancy results from a combination of the low-level implementation errors detailed in the supplementary text and a more conceptual error specific to the design of this analysis – the authors balanced trials for choice on a per stimulus basis when training their decoder, which amounts to training on flat psychometric curves, effectively baking into the decoder an insensitivity to features that correlate with choice (such as intensity).

Altogether, our findings demonstrate that DLS dynamics and animals’ duration judgments were similarly modulated by the intensity of the second stimulus.

### DLS neural responses displayed negligible sensitivity to the intensity of the first stimulus

We observed that the intensity of the first stimulus had no systematic net effect on animals’ duration judgments (Fig. 2C, F, S2). As such, the possibility that this stimulus feature affected DLS activity past S1 and the ISI and into the S2 period would be hard to reconcile with the DLS being a substrate for said I1-independent decisions. To address this, and for ease of comparison, we applied the exact same treatment used to characterize the impact of I2 on S2-aligned DLS activity in the previous section (Fig. 3) to I1-conditioned activity recorded during S2 (Fig. 4). We did not observe any orderly relationship between I1 and S2-aligned: population averages (Fig. 4A); single-neuron responses (Fig. 4B, same examples from Fig. 3B); PC projections (Fig. 4C); PC trajectories (Fig. 4D); neurometric curves (Fig. 4E); and decoded time estimates (Fig. 4F).

**Figure 4:**
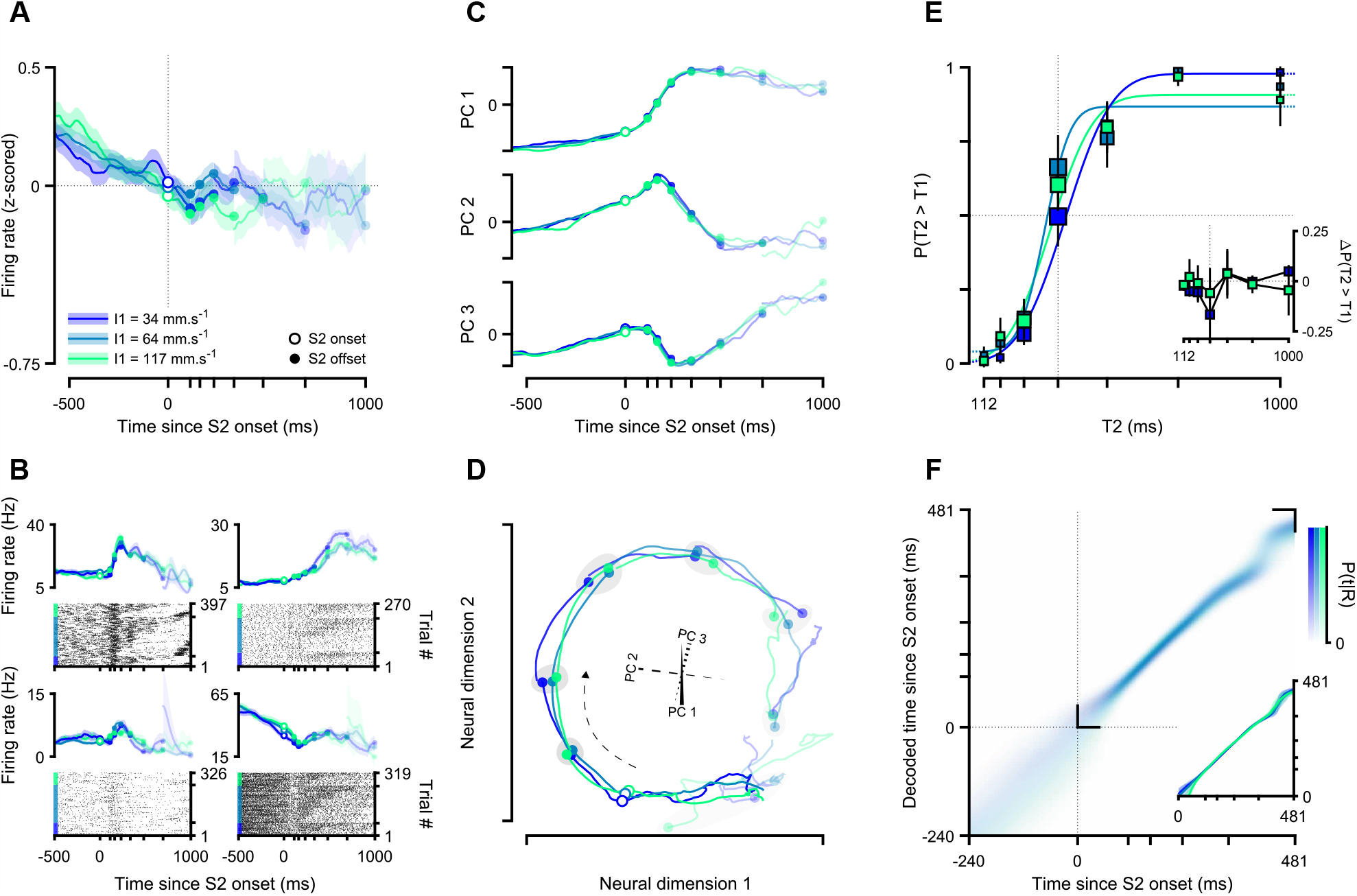
DLS dynamics recorded during the second stimulus were unaffected by the intensity of the first stimulus. (**A**) Average normalized population activity conditioned on I1 and aligned to the onset of S2 (mean ± SEM across neurons). (**B**) Responses of four example DLS neurons during S2. For each neuron, spike density functions (mean ± SEM across trials) are conditioned on I1 and plotted above its spike raster, where trials are first sorted by I1, and then by T2 within each I1 level. (**C**) I1-split population activity projected onto each of the first three PCs computed using I1-agnostic population activity aligned to the onset of S2. (**D**) Population activity conditioned on I1 projected onto a 2D section of the subspace defined by the three PCs shown in (C). The 3-axis gizmo at the center depicts the projections of the axes defined by the first three PCs onto the chosen 2D section, the dashed black arrow indicates the direction of time, and the gray patches group population states recorded at each possible stimulus offset across I1. (**E**) Neurometric curves trained using S2-aligned population activity from middle-I1 trials conditioned on I1 (main axes), and differences between the proportion of T2 > T1 neural “reports” in high (I1 = 117 mm s^−1^) and low (I1 = 34 mm s^−1^) intensity trials compared to middle intensity (I1 = 64 mm s^−1^) trials (inset, mean ± propagated SEM). (**F**) Time estimates decoded from DLS activity around S2 presentation conditioned on I1. Main axes: mean posterior distributions across runs. Inset: MAP point estimates averaged across runs (median and IQR). The opacity of lines, markers and patches in (A-D) reflects the relative number of trials contributing to the underlying estimates.

Furthermore, I1 modulation of DLS dynamics during the S1 period was also virtually non-existent, if not entirely absent (Fig. S4), thus standing in stark contrast to the effect of I2 on S2-aligned activity (Fig. 3). Since there was nothing experimentally different about the delivery of the two stimuli, this striking asymmetry argues against the DLS faithfully encoding all aspects of an animal’s immediate sensory state, and in favor of its activity reflecting a learned mapping between states and actions. It should be noted that the authors reported this uneven effect of intensity on DLS activity during the two stimuli in their erratum (Toso, Reinartz, et al., 2021), though their qualitatively correct observation was unsatisfactorily accounted for in the absence of the accompanying characterization of the asymmetric impact of I1 and I2 on behavior.

Taken together, these data speak to a remarkable indifference of DLS responses to the intensity of the first stimulus relative to that of the second.

### The speed of DLS dynamics covaried with duration judgments

Systematic correlations between the speed of striatal dynamics and interval timing have been observed across laboratories, animal models, interval timing paradigms and striatal domains (Gouvêa et al., 2015; Kim et al., 2018; Mello et al., 2015; Monteiro et al., 2023; Wang et al., 2017). Adding to the evidence provided by these correlative studies, our group has recently published evidence in support of this link being causal, by using temperature to slow down and speed up population dynamics in the dorsal striatum of rats and observing corresponding biases towards short and long duration judgments (Monteiro et al., 2023). Perhaps then the most surprising of the central claims of Toso and colleagues was that there was no relationship between DLS activity and animals’ duration discrimination. Importantly, this was inferred from the overlap in time estimates decoded from S1- and S2-aligned DLS activity recorded during correct and incorrect trials. This analysis is problematic not just because nominal correctness can not be used to meaningfully dichotomize rats’ behavior (Fig. 1) and because rats mostly used S2 to guide choice (Fig. 2), but more importantly because, even assuming rats performed as idealized, of how correct and incorrect trials map onto over- and underestimation of duration changes across time within each stimulus period. Incorrectly discriminating stimulus pairs for which T2 is longer than T1 presumably corresponds to faster-than-usual timekeeping during S1 and/or slower-than-usual timekeeping during S2, whereas the opposite is true for errors when T2 is shorter than T1 (Fig. S5). The same logic holds in reverse for correct trials. Therefore, pooling decoder output conditioned on trial outcome indiscriminately across stimulus pairs would obscure these signatures within each category for all but extreme T1 and T2 values (equal to or longer than 694 ms), since these durations signal which choice is correct unambiguously and do not overlap in time with any other (ambiguous) durations. However, since the authors had to exclude these trials due to data scarcity, their analysis was incapable of detecting even this remnant of the correlation between time as encoded by the DLS and rats’ choices.

To circumvent this issue, we instead conditioned neural activity on whether rats judged T2 to be greater than T1, which should partition trials into two groups with consistent opposite underlying biases in timekeeping during the presentation of S2. We thus subjected S2-aligned DLS activity conditioned on choice to the same battery of analyzes we used for I2 (Fig. 3) and I1 (Fig. 4, S4). Consistent with previous findings, we saw that DLS responses covaried with animals’ judgments. Interestingly, choice modulation, though present, was perhaps not as evident as that driven by I2 when looking at population mean activity (Fig. 5A) or responses from the individual neurons we have been showcasing throughout (Fig. 5B). The reason for this became apparent with PCA (Fig. 5C, D), which revealed that unlike intensity-, decision-related variability seems to express itself almost exclusively through the speed of DLS dynamics. This was corroborated by both the horizontal shift in neurometric curves (Fig. 5E) and the steady divergence between decoded time estimates conditioned on animals’ reports (Fig. 5F), which can be understood as the accumulating effects of slow-biased (fast-biased) neural trajectories preceding T2 < T1 (T2 > T1) reports resulting in population states at S2 offset that increasingly lag behind (lead ahead) their T2 > T1 (T2 < T1) counterparts.

**Figure 5:**
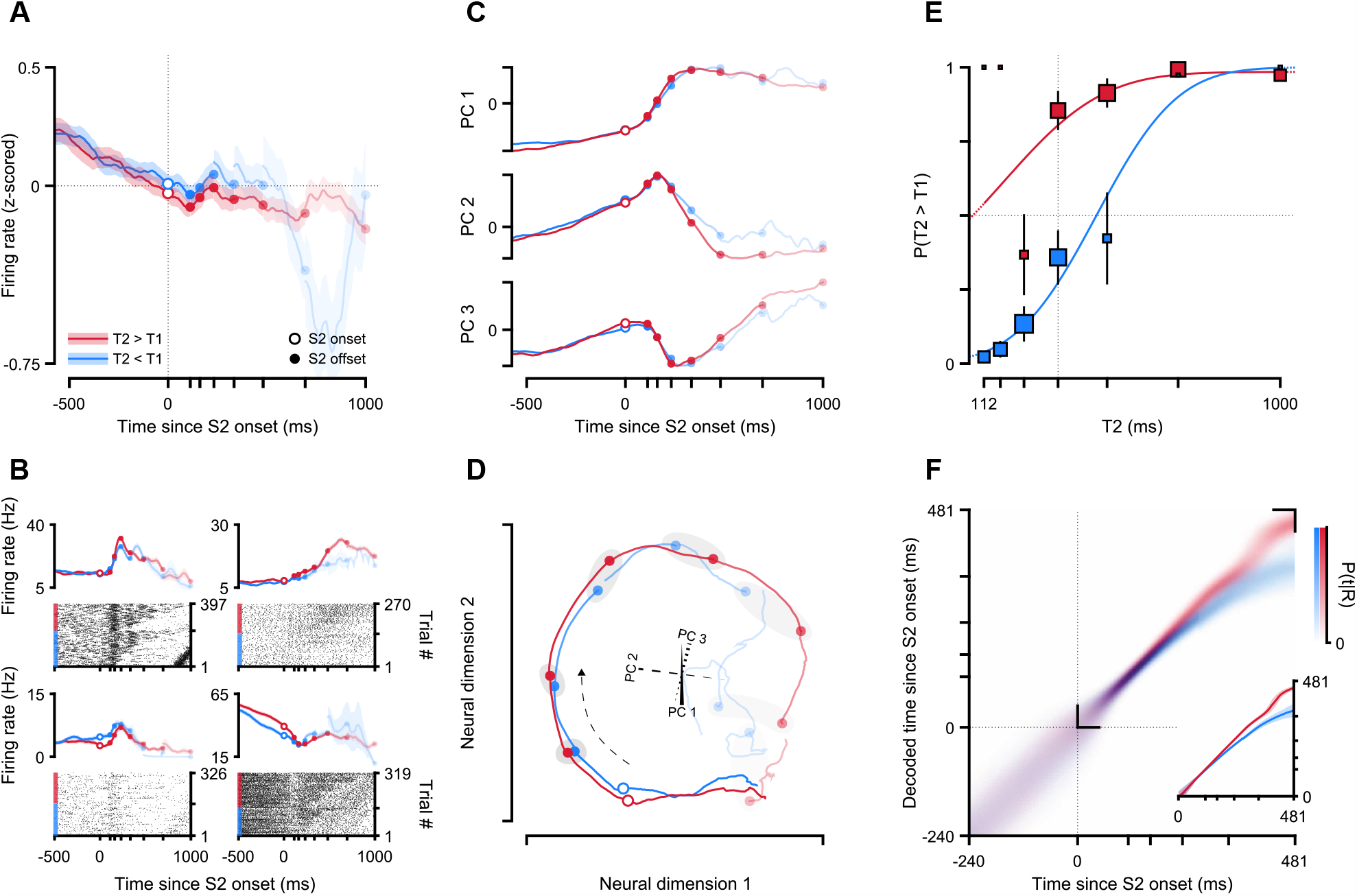
The speed of DLS dynamics covaried with duration judgments. (**A**) Average normalized population activity conditioned on choice and aligned to the onset of S2 (mean ± SEM across neurons). (**B**) Responses of four example DLS neurons during S2. For each neuron, spike density functions (mean ± SEM across trials) are conditioned on choice and plotted above its spike raster, where trials are first sorted by choice, and then by T2 within each choice category. (**C**) Choice-split population activity projected onto each of the first three PCs computed using choice-agnostic population activity aligned to the onset of S2. (**D**) Population activity conditioned on choice projected onto a 2D section of the subspace defined by the three PCs shown in (C). The 3-axis gizmo at the center depicts the projections of the axes defined by the first three PCs onto the chosen 2D section, the dashed black arrow indicates the direction of time, and the gray patches group population states recorded at each possible stimulus offset across choice categories. (**E**) Neurometric curves trained using S2-aligned population activity from choice-agnostic trials conditioned on choice. (**F**) Time estimates decoded from DLS activity around S2 presentation conditioned on choice. Main axes: mean posterior distributions across runs. Inset: MAP point estimates averaged across runs (median and IQR). The opacity of lines, markers and patches in (A-D) reflects the relative number of trials contributing to the underlying estimates.

Together these results demonstrate that judgments about the duration of the second stimulus could be reliably predicted by the speed with which patterns of DLS activity evolved during its presentation.

### The *relative* proportion of DLS neurons modulated by different task variables matched their relative impact on performance

A surprising difference between our analyzes and those presented in the original paper and its correction is in the number of neurons identified as modulated by stimulus intensity. We found this discrepancy to be driven by a combination of issues related to the particular statistical criteria the authors enforced.

It is increasingly appreciated that single neuron activity is often modulated by multiple factors (Kobak et al., 2016; Mante et al., 2013; Rigotti et al., 2013), indicating that to fully understand a neuron’s response properties, both sound experimental design that unconfounds task variables and analysis methods that quantify their relative contributions are necessary. In this section we provide a summary for how different task variables affected the activity of DLS neurons at different moments in the task, ultimately more focused on I2 during S2 to explore a technical issue regarding the operational definition of intensity-selective neurons used by Toso and colleagues – for a neuron to be considered as modulated by intensity, spike counts aligned to the onset of either stimulus were required to be significantly correlated with intensity (of that stimulus) for at least five consecutive and non-overlapping 100-ms bins. We find that these criteria are statistically underpowered to detect cells modulated by intensity in light of the typical firing statistics of striatal neurons (Steiner & Tseng, 2016) and the total amount of data available. The nature of this task (and most other sensory timing tasks) is such that the number of trials recorded during the presentation of a given stimulus decreases at discrete boundaries defined by the set of its possible durations. This means that with the authors’ approach, statistical power drops sharply with each subsequent regression, to the point where only trials corresponding to the two longer stimulus durations (694 and 1000 ms) can contribute past the fourth bin-wise regression. Yet, all regressions were treated equally in how they contributed to a neuron being labeled as modulated by intensity. Furthermore, considering the characteristically low firing rates of medium spiny neurons (MSNs, the vast majority of striatal neurons, Steiner and Tseng, 2016), the response variable – spike counts recorded within 100 ms – is effectively varying between 0 and 2, thus impairing statistical power even further.

To try and navigate these constraints while facilitating a direct comparison with the authors’ approach, we fit separate linear regression models to each neuron’s spike counts aligned to multiple task events (trial initiation, S1, ISI, S2 and “go” cue), and calculated the proportions of neurons that were significantly up- and down-modulated by the different regressors (Fig. 6A). Crucially, instead of regressing spike counts against I1 or I2 separately, we used a multivariable design matrix that included not just I1 and I2, but also T1 and T2 as well as choices and rewards in the current and previous trials. This was done so as to account for any systematic (Fig. 1C, F) as well as spurious correlations between predictors, and thus better parse variance that reflected direct modulation by each stimulus feature independently of one another as well as of premotor-, motor-, and reward-related signals.

**Figure 6:**
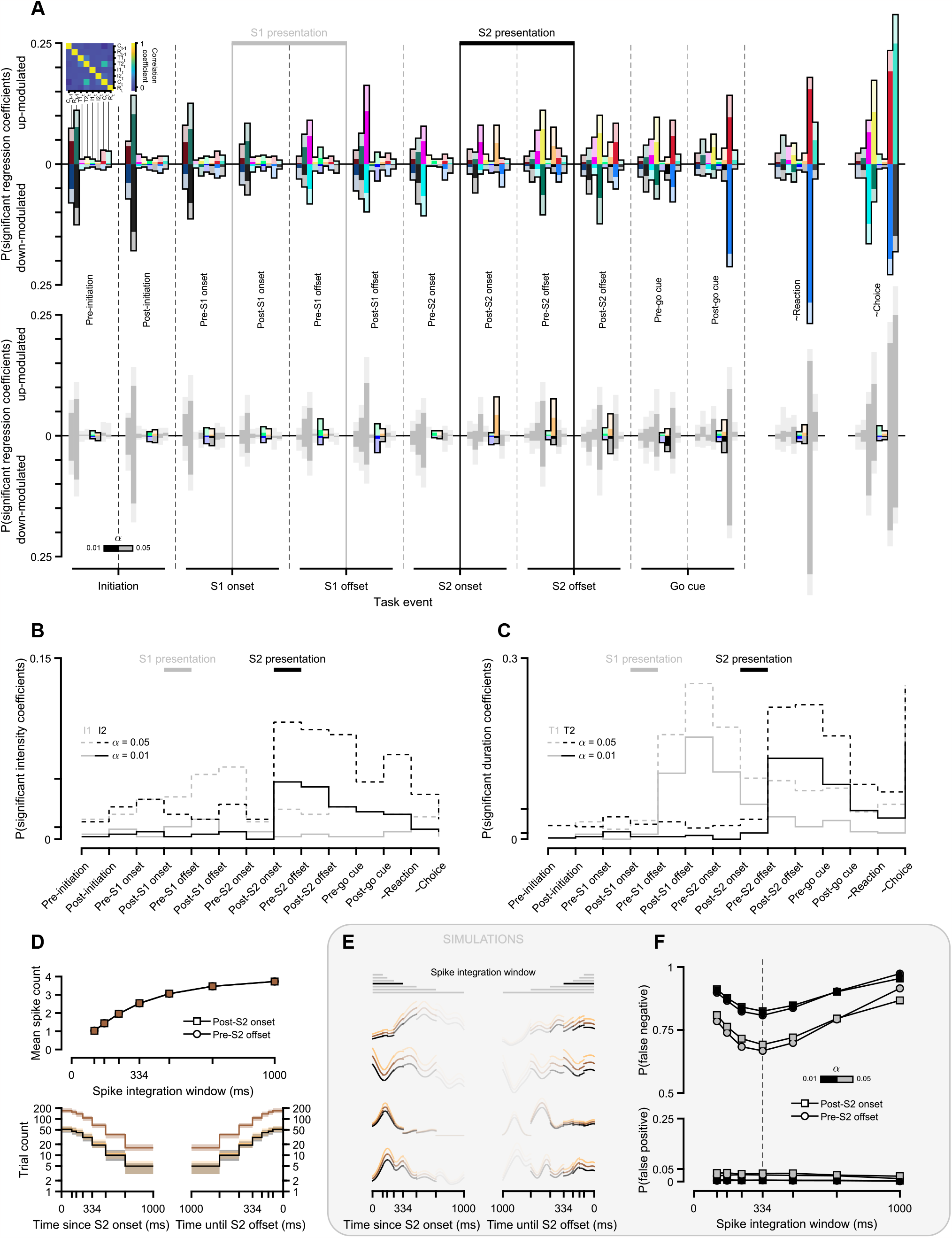
The *relative* proportion of DLS neurons modulated by different task variables matched their relative impact on performance. (**A**) Top: proportion of DLS neurons that were either up- or down-modulated by several experimental and behavioral factors across multiple task epochs. Signed modulation was assessed by quantifying the proportion of significant positive and negative coefficients in multivariable linear regression models fit to spike counts recorded in a 334-ms window aligned to the onset and/or offset of multiple task events. The “∼” in ∼reaction and ∼choice refers to the approximate nature of the alignment for these epochs (334 ms and 668 ms after go cue for reaction and choice, respectively), which lacked trial-to-trial annotated event times. The factor/regressor cross-correlation matrix is shown as an inset in the top left. Bar colors are factor-specific and respect all previously introduced color schemes (where applicable), and bar opacity refers to the significance level used to assess modulation (1% and 5% for opaque and transparent colors, respectively). Bottom: same as above, but highlighting modulation by I1 and I2. (**B**) Total proportion of DLS neurons that were up- or down-modulated by I1 (in gray) and I2 (in black) across the different task epochs. (**C**) Same as (B), but for stimulus durations. (**D**) Opposing effects of the duration of the spike integration window used in (A-C) on the average number of recorded spikes and the number of trials contributing to that estimate during S2 presentation epochs. Top: average number of spike counts across trials as a function of spike integration window for the intermediate-I2 condition. Bottom: number of trials with recorded spike counts at each time point relative to the onset (left) or offset (right) of the second stimulus, split by I2 condition (median and IQR across neurons). (**E**) I2-conditioned generative spike density functions used in simulated positive controls for the same four example DLS neurons shown in previous figures, aligned to the onset (left) or offset (right) of the second stimulus. (**F**) False negative (top) and false positive (bottom) rates computed by calculating the proportion of I2-modulated neurons during the S2 presentation epochs averaged over two separate sets of 25 runs corresponding to positive and negative controls. For each run 100% of the simulated neuronal responses were either positively modulated by I2 (when computing false negative rates) or not modulated by I2 (when computing false positive rates).

As mentioned above, in the specific case of stimulus epochs, the fact that most neurons were only recorded for a few trials for any given combination of vibration durations and intensities means that as time relative to the onset (offset) of the stimulus increases (decreases), the number of trials that continue to meet the selection criteria up to (from) that point decreases. This leads to a trade off in the selection of the *spike integration window* for this analysis. Longer (shorter) windows will lead to a larger (smaller) range in the support of the distribution of spike counts over trials (Fig. 6D, up), but at the cost of fewer (more) trials contributing samples to that distribution (Fig. 6D, bottom). To both inform the choice of this parameter and create negative and positive controls for this analysis, we generated surrogate spike data with similar firing statistics to those found in the real data. Briefly, for each neuron we first computed the cross-trial average spike density function aligned to the onset of S2 using only intermediate-I2 trials (I2 = 64 mm s^−1^). This average response was then either used directly as the generative spike rate function for all trials in the case of negative controls, or first multiplied by a gain parameter proportional to I2 in the case of positive controls (Fig. 6E). The resulting rate functions were then used to sample spike times from inhomogeneous Poisson point processes on a trial-by-trial basis. Running our regression analysis with these surrogate data allowed us to estimate false positive (type I errors) and negative rates (type II errors) for a range of spike integration windows, which suggested 334 ms as the optimal time window given the previously mentioned constraints imposed by data scarcity (Fig. 6D). Even with this optimized window, the estimated false negative rate was close to 70% (80%) at a significance threshold of 5% (1%) (Fig. 6F), meaning that our detected 9.7% (4.7%) of I2-modulated neurons (Fig. 6B, left) may actually reflect that as much as 32.3% (23.5%) of DLS neurons were modulated by the intensity of the second stimulus during its presentation.

Granted, these error rates are intimately tied to the nature and magnitude of intensity modulation we simulated in our positive controls, which, though tuned to approximately match effect sizes seen in the data (compare Fig. 3B and Fig. 6E), were not explored systematically. Still, it is worth stressing that for simplicity, in our positive controls intensity had a multiplicative effect on the amplitude, and not the temporal scale, of generative spike rates. Modulation through temporal scaling would arguably be even harder to detect, due to the need for its effects to accumulate over time, and how that interacts with the number of contributing trials decreasing as a function of elapsed time since stimulus onset (Fig. 6D, bottom). Case in point, the proportion of neurons modulated by the upcoming choice, which we have shown to manifest almost entirely in the speed of population dynamics (Fig. 5), seem comparatively low during S2 (Fig. 6A). This means that the estimated false negative rates may well be underestimating the true rate of type II errors in our quantification. For all of these reasons, instead of reporting an “exact” fraction of intensity-modulated neurons, that would be inextricably linked to arbitrary parameters in its calculation (e.g., a spike integration window lasting *x* ms and a significance level of *y*), cell type-specific firing statistics, and what features of their responses were affected, we emphasize that it was definitely not zero, nor was it two percent.

Regarding modulation by other task variables, such as I1, choice and reward, we again find that the authors’ estimates were systematically below ours (even before accounting for false negative rates, which would have to be estimated separately on an epoch-by-epoch basis), but would once more caution against statements about the absolute fraction of neurons displaying sensitivity to a particular task factor. However, *relative* statements strike us as potentially meaningful, depending on the question at hand. For example, intensity modulation was much more apparent during and after the second stimulus than during and after the first, where it was comparatively absent (Fig. 6A, B). In yet another example, DLS activity strongly reflected the duration of S1 early in the ISI period, but the proportion of neurons carrying information about which T1 had been presented seemed to steadily decrease across subsequent epochs, to the point where it was roughly between one third and one sixth (depending on significance thresholds) the proportion of neurons modulated by T2 right after the offset of S2 (Fig. 6A, C). This ratio of neural modulation is reminiscent of the ratio of contribution of these variables onto rats’ decisions (Fig. 2C), and is consistent with previous proposals that contraction bias arises not at the encoding stage (during S1), but gradually up to the decision stage (presumably during or shortly after S2, Ashourian and Loewenstein, 2011).

Together, these results reiterate a fundamental aspect of the data – I2 modulation was prevalent among DLS neurons – and moreover, they underscore the accordance between the extent to which different stimulus features affected temporal judgments and modulated neural activity across the DLS.

### Differences in time decodability from putative clusters of ramping and non-ramping DLS neurons were biased by confounds unrelated to those category labels that were introduced by categorization criteria

Two of the seven main figures in Toso, Reinartz, et al., 2021 were devoted to the issue of *ramping* vs. *non-ramping* neurons in the DLS, and how time estimates decoded from ramps were more reliable than those decoded from non-ramps. While we understand in theory the interest in ramping profiles due to long-existing computational accounts of timing as resulting from an integration process (Gibbon, 1991; Treisman, 1963), their existence as a distinct functional class within neural populations is by no means axiomatic. Yet, like others before them (Donnelly et al., 2015; Emmons et al., 2017), the authors assumed these categories exist and applied several ad-hoc criteria to classify neurons into the two groups without first providing evidence for the clusterability of their activity profiles. Additionally, though the differential time encoding of putative ramps and non-ramps was ultimately only assessed during stimulus presentation periods, these were not the epochs used in the clustering steps. Instead, the authors first enforced their classification criteria in two 1-s windows centered on the onset and offset of each stimulus, and later labeled the union of up- and down-ramping neurons identified in those stimulus-flanking windows as stimulus period ramps (and all remaining neurons as non ramps, Fig. 7A, bottom). In other words, a neuron was considered to ramp during the stimulus period if it ramped (up- or down-) from 500 ms before to 500 ms after stimulus onset and/or offset, leading to an odd set of possibilities: a neuron ramping exclusively before or exclusively after stimulus presentation will be labeled a stimulus *period* ramp, regardless of its response profile during stimulus presentation; a neuron with a unimodal response profile (a “bump”) peaking half way through a given stimulus period will be considered a ramp (since its activity increases going forward from stimulus onset and decreases going backward from its offset). Evidence for this last scenario at play can be seen in the contrasting shapes of the distributions of temporal tuning (see methods) of ramps during the task epochs in which these were first identified – bimodal – and during stimulus epochs – unimodal (Fig. 7A top). In all cases, these were statistically different from the strictly unimodal temporal tuning distributions of non-ramps, even during stimulus epochs (Fig. 7A top).

**Figure 7:**
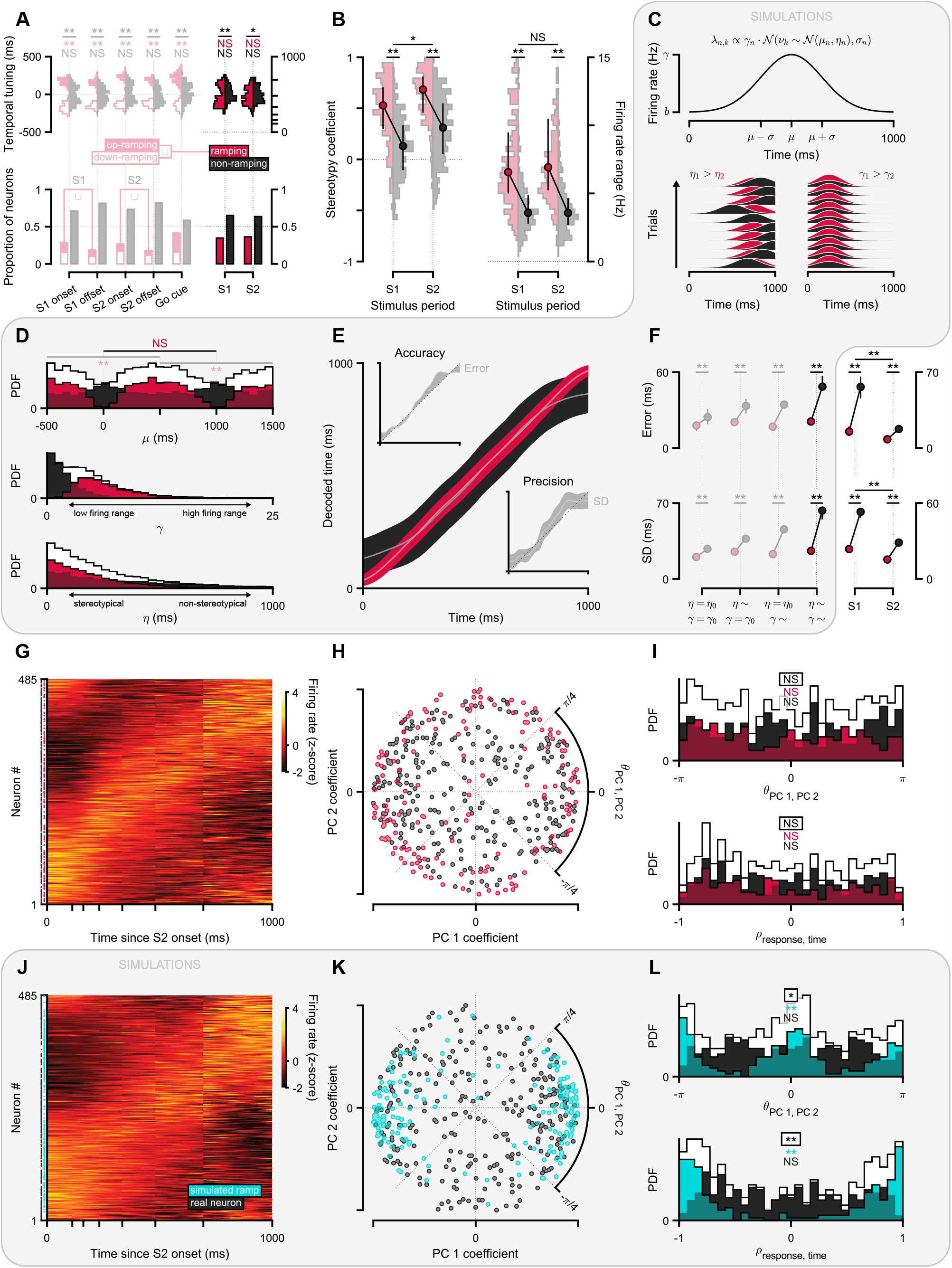
Criteria for categorizing neurons within a continuum can artifactually influence decoded time estimates. (**A**) Bottom: proportion of DLS neurons labeled as ramping (up and down, where applicable) and non-ramping using the classification and grouping criteria from Toso and colleagues. Task epochs in which classification criteria were enforced are washed out, and stimulus epochs used to assess decoding performance for each cluster are vivid. Ramps in each stimulus period (S1 and S2) do not preserve their initial direction qualifier because they were defined as the union of up- and down-ramping labels across the respective onset and offset epochs (an S2 ramp might have ramped up during the S2 onset epoch and down during S2 offset epoch). Top: cluster-split distributions of temporal tuning across DLS responses recorded during 1-s windows centered on each task event (washed out), or aligned to the onset of each stimulus period (vivid). (**B**) Distributions of stereotypy coefficients (left) and firing rate ranges (right) conditioned on the authors’ cluster identities. (**C**) Specification of the model used to simulate Gaussian bump responses (top) and generative spike rates in ten example trials of two pairs of simulated neurons that varied either in their stereotypy (bottom left) or gain (bottom right) parameters. (**D**) Distributions of temporal tuning (top), stereotypy (middle) and gain (bottom) parameters agnostic to (in white) or split by whether the corresponding simulated neurons were labeled as ramps or non-ramps using the authors’ criteria. Horizontal gray and black bars at the top represent the epochs in which Toso et al. enforced their clustering criteria and evaluated decoding performance, respectively. (**E**) Main axes: decoded time estimates from two equally sized subpopulations (N = 150) of Gaussian bumps varying along their stereotypy, gain, and temporal tuning parameters, and conditioned on whether they were ramps or non-ramps (mean posterior mean ± mean posterior SD across 100 simulations, each with 300 trials). Insets: decoded time estimates from a single simulated trial (posterior mean ± posterior SD) illustrating the decoding accuracy (top) and precision (bottom) metrics used in (F). (**F**) Comparison of decoding accuracy (top) and precision (bottom) from ramping and non-ramping neurons, for simulations from four differently parameterized models (left) and DLS data recorded during S1 and S2 (right, median and IQR across simulations or bootstrap iterations). Model variants were defined by whether their stereotypy and/or gain parameters were fixed (*η* = *η*_0_ and/or *γ* = *γ*_0_) or sampled (*η* ∼ and/or *γ* ∼) from the corresponding distributions shown in white in (D). (**G**) Normalized spike density functions of all DLS neurons recorded during S2. Neurons are ordered by polar angle in the subspace defined by the first two PCs describing firing dynamics during S2. (**H**) Joint distribution of PC coefficients conditioned on the authors’ cluster identities. (**I**) Distributions of clusterability metrics – polar angle of PC coefficients shown in (H) (top) and Pearson’s correlation coefficient between the responses shown in (G) and time (bottom) – either agnostic to (in white) or split by cluster identity. (**J-L**) Same as (G-I), but for a hybrid simulation wherein neurons labeled as ramps using the authors’ criteria were replaced by simulated ramps (evenly split between up- and down-ramping).

In addition to the issue of *when* classification criteria were applied, we found the criteria themselves problematic considering that the purpose of this analysis was to assess differences in time decodability from activities in the two resulting classes. First, to be classified as a ramp, a neuron’s firing rate was required to be stereotyped over trials. Were this criterion used to select cells for further analysis in general, we see no issue. However, non-stereotypical cells were instead placed in the non-ramping category. Unsurprisingly, this produced a marked asymmetry in the stereotypy coefficients (see methods) of ramps and non-ramps during both S1 and S2 (Fig. 7B, left). It then becomes circular that decoding performance, which hinges on the reproducibility of neural responses across trials, was better with ramping than with non-ramping neurons. Second, as a proxy for a monotonicity requirement, the average firing rate of ramping neurons was required to be significantly correlated with elapsed time, and with a linear regression slope that crossed an arbitrary threshold in *absolute* value. This last detail is again problematic, as it selects for neurons with high dynamic range in their firing rate over those with a relatively lower range (Fig. 7B, right). Intuitively, this translates into systematically poorer temporal informativeness in non-ramps (just as an image with an 8-bit color depth can resolve fewer colors than a 24-bit one). Additionally, because firing rate ranges correlate with overall levels of firing rate (data not shown), lower ranges also translate into higher variability in spike timing, which further hinders (benefits) precision when decoding time from the responses of non-ramps (ramps).

To validate these intuitions, we rigorously applied the authors’ operational definitions of ramping and non-ramping neurons to a population of simulated Gaussian bumps with variable temporal tuning (when they peak on average), stereotypy (how much peak times vary across trials), and gain (how high are those peaks relative to baseline), each controlled by independently sampled parameters specific to each simulated neuron (Fig. 7C).

As expected, simulated bumps labeled as ramps were systematically associated with higher stereotypy and gain parameters, as well as with distributions in temporal tuning parameters that were bimodal in flanking epochs (in which classification occurred) and unimodal in the epoch of interest (in which decoding performance was evaluated, Fig. 7D). Despite this difference in shape, and similarly to what was observed in DLS responses, temporal tuning distributions during the decoding epoch were different for the two groups (Fig. 7A, D).

Altogether, these selection biases greatly impaired the accuracy and precision of time decoding in the non-ramping category relative to the ramping one (Fig. 7E, F, left), though they were each sufficient individually to produce significant changes in decoding performance (Fig. 7F, left). This exercise demonstrates how the authors’ results, which we also replicate here (Fig. 7F, right), derive from artifacts induced by their selection criteria unrelated to the original objective of identifying neurons with different temporal response profiles. Put differently, the criteria pre-determined the result. Incidentally, it bears noting that in line with the asymmetric contribution of S1 and S2 onto rats’ decisions, decoding performance (agnostic to cluster identities) was significantly better during S2 than S1 (Fig. 7F, right), likely due to (at least in part) more stereotypical DLS responses during the second stimulus relative to the first (Fig. 7B, left).

In summary, these results demonstrate that the authors’ assertion regarding the relatively superior decodability of elapsed time from ramping as compared to non-ramping DLS responses is unfounded.

### No evidence for the existence of ramping and non-ramping clusters in DLS responses recorded during the second stimulus

We show above how the authors’ categorization criteria for labeling a neuron as ramping or non-ramping was problematic for the validity of claims regarding relative accuracy of time decoding in the two categories of cells. However, we wondered whether there was any structure in the data that would suggest such categories in the first place. If the data is deemed to not possess sufficient structure to be meaningfully partitioned, then clustering becomes arbitrary, potentially misleading and arguably unwarranted (Adolfsson et al., 2019). We thus sought to assess in an unbiased manner whether there is evidence for categories of DLS response profiles in the data, and, if so, whether those categories correspond to ramping and non-ramping patterns of activity. To do this, we adopted the *clusterability via multimodality approach*, which formalizes the intuitive notion that if clusters do exist, distributions along features that define them must be multimodal (Adolfsson et al., 2019).

To evaluate our ability to detect ramps should these exist, we analyzed the clusterability of two data sets. A *real* one, consisting of DLS activity recorded during the presentation of the second stimulus (Fig. 7G-I), and a *hybrid* one that consisted of those very same S2-aligned real responses but where the neurons originally labeled as ramps (using the criteria from Toso et al.) were replaced with actual simulated ramps (evenly split between up- and down-ramping, see methods, Fig. 7J-L). As a starting point, in both cases we plotted normalized spike density functions for all neurons ordered according to their temporal profile of activity (Fig. 7G, J). Specifically, responses were sorted by the angle of their first two PC coefficients in polar coordinates. We did not visually detect discontinuities in firing profiles themselves, which appeared more like a continuum. Although this was true even in the hybrid case, where we know there were ramps, methods used to order cells are known to produce visually misleading effects. Further inspection of the joint distribution of the PC coefficients used to sort those neural response rasters revealed that indeed simulated ramps tended to cluster in two poles opposite to each other, corresponding to up- and down-ramping neurons (Fig. 7K). The same phenotype was not observed in the case of ramps inferred by the authors among strictly real DLS activities, which instead seemed much more spread out in PC space, if not for displaying a slight tendency towards higher eccentricities (Fig. 7H). This bias along the radial polar coordinate was to be expected given the previously described bias in firing rate ranges induced by the clustering method (Fig. 7B, right).

To quantify these observations and formally assess the existence of ramping and non-ramping clusters, we selected two features that satisfied the recently proposed *efficiency, algorithm independence* and *effectiveness* requirements for well-suited measures of clusterability (Adolfsson et al., 2019): 1) polar angle in PC coefficient space, and 2) Pearson’s correlation coefficient between response profiles and elapsed time since the onset of S2. For both measures, we found that their distributions were unimodal in the case of real data (Fig. 7I), and multimodal in the case of hybrid data (Fig. 7L). Furthermore, real data distributions restricted to putative ramps remained unimodal, indicating that the authors’ classification approach fell short of even dichotomizing a continuum (Fig. 7I).

These results argue strongly against DLS activity profiles naturally clustering into ramping and non-ramping subpopulations, and instead favor a more holistic view of the dynamics underlying time encoding by populations of striatal neurons.

## Discussion

We have shown that the activity of neural populations in dorsal striatum correlates with animals’ judgments of stimulus duration, consistent with a growing number of studies that have focused on a variety of different behavioral tasks (Bakhurin et al., 2017; Gouvêa et al., 2015; Jin et al., 2009; Kim et al., 2018; Matell et al., 2003; Mello et al., 2015; Monteiro et al., 2023; Wang et al., 2017). Despite our contrasting conclusions to those of the original authors, we find the data set to be a valuable addition to this body of work, in that the behavioral task leverages an innate tendency for duration judgments to be biased by stimulus intensity to study potential neural correlates of timing behavior. Prior work employing motor timing tasks has involved training animals to respond at different times, depending on the identity of an explicit cue or recent trial history with respect to temporal availability of rewards (Malapani & Fairhurst, 2002; Mello et al., 2015). Temporal judgment tasks, or what are sometimes referred to as sensory timing tasks, often leverage trial by trial variability in judgments applied to particular durations to ask similar questions (Paton & Buonomano, 2018). However, the behavioral task used here takes advantage of the fact that stimuli of differing intensity are subject to systematically modified temporal estimation processes to take experimental control over internal mechanisms related to the judgment of duration (Toso, Fassihi, et al., 2021; Toso, Reinartz, et al., 2021). The fact that modification of temporal judgments via this apparently innate mechanism caused similar changes in striatal activity to those observed when timing behavior is modified using other means further strengthens the hypothesis that patterns of striatal activity indeed underlie animals’ temporal judgments.

How does this relate to the notion of temporal “perception” on which the original study focused to a large degree? We remain agnostic to the animal’s subjective experience, and thus prefer the term “judgment” which we operationally define as the animal’s report. To perform this task, animals must transform information regarding the duration of vibratory sensory stimuli into a binary report, or choice, but also obey other requirements of the task, such as initiating a trial and maintaining their presence in a nose port for an extended period (Cruz et al., 2022). All of these elements of the behavior involve the mapping together of sensory or internal variables with particular actions, and implicitly require a time-base as part of that process. In other words, during training animals learn a policy for behavioral control that must rest on a temporal basis. A wealth of data implicate the striatum in learning policies for behavioral control that maximize expected reward (Cruz et al., 2022; Lau & Glimcher, 2008; Reynolds et al., 2001; Samejima et al., 2005; Schultz et al., 1997; Soares et al., 2016; Tai et al., 2012). Thus, we tend to view striatal correlates of timing behavior as reflecting how neural dynamics provide a temporal basis for constructing intermediate representations required for behavioral control policies, such as the value of state-action combinations, which may or may not impinge on the animals’ subjective experience of duration. That said, the animal’s pattern of choices is the only window we have as external observers into any potential subjective percept of interval duration that may exist. The correspondence between timing behavior and neural activity in the striatum, reproducibly observed across many studies with differing behavioral task designs, suggests that striatal activity may at least reflect such a putative subjective perception of duration, independent of its ultimate origin.

## Acknowledgements

We thank T. S. Duarte, M. Sousa and S. A. Zamora for comments on versions of the manuscript and the entire Paton laboratory, past and present, for feedback during the course of this project. The work was funded by an HHMI International Research Scholar Award to J. J. P. (55008745), a European Research Council Consolidator grant (DYCOCIRC-REP-772339-1) to J. J. P., a Bial bursary for scientific research to J. J. P. (193/2016), internal support from the Champalimaud Foundation, and a PhD fellowship from the Portuguese Foundation for Science and Technology (FCT) to F. S. R. (SFRH/BD/130037/2017). T. M. was supported by a grant from FCT (CDL-CTTRI-249-SGRH/2022) and through multiannual funding to the WJCR (UIDB/04810/2020). We thank the support of NVIDIA Corporation with the donation of the Titan X Pascal GPU used for this research. The funders had no role in study design, data analysis, decision to publish or preparation of the manuscript.

## Author contributions

F. S. R. analyzed the data and drafted and edited the manuscript. T. M. and A. M. reviewed and edited the manuscript. J. J. P. supervised all aspects of the project and drafted and edited the manuscript.

## Data availability

Behavioral and neural data were made available by Toso, Reinartz, et al., 2021 at: https://doi.org/10.17632/wp9h39kbtv.2.

## Code availability

Analysis code that supports the findings of the original and the present studies are respectively available at:

https://doi.org/10.17632/wp9h39kbtv.2

https://github.com/flipe-rodrigues/toso2021-comment

## Methods

### Behavioral analyzes

#### Psychometric functions

We used the *Psignifit* toolbox (Schütt et al., 2016) to fit the following 4-parameter psychometric function to all duration discrimination data:

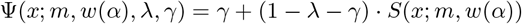

In this parameterization, a strictly monotonic sigmoid function *S* from the stimulus level *x* (corresponding to either T1 or T2) onto the unit interval [0, 1] is specified by *m* = *S*^−1^(0.5) and *w* = *S*^−1^(1 − *α*) − *S*^−1^(*α*), namely the *threshold* and *width* parameters. This is independent of the choice of *S*, which in our case was the reversed cumulative Gumbel distribution. The hyper-parameter *α*, which sets the span of *w* along the vertical axis, was set to 0.05. To account for stimulus-independent choices, *S* is scaled by two additional free parameters, *λ* and *γ*, which respectively control the upper and lower asymptotes of the psychometric function Ψ.

#### Psychometric model comparison

To assess whether and how stimulus intensity impacted specific parameters of the psychometric function described above, we first built a reference psychometric curve for each stimulus (S1 and S2), by fitting all of its four parameters to rats’ choices as a function of stimulus duration using only middle-intensity trials. Then, for each of the remaining intensity conditions (high and low), we fit a battery of models that varied combinatorially in which parameters were let free and which were fixed to those of the corresponding middle-intensity reference model. Lastly, we computed the Bayesian information criterion (BIC) for each tested model and each tested intensity condition, and took relatively lower (higher) BIC scores as evidence for (against) the corresponding models.

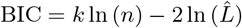

Where *n* is the number of observations (trials), *k* is the model-specific number of free parameters and 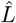 is the corresponding maximum likelihood.

### Single neuron analyzes

#### Spike density functions

Spike density functions (spike density functions (SDFs)) were built on a trial-by-trial basis by taking the provided histograms of spike times (computed by the original authors using a 1-ms bin) and convolving them with a causal kernel specified by a gamma distribution with shape and scale parameters of *k* = 2 and *θ* = 50 ms, respectively. Normalized SDFs were computed as z-scores of stimulus-aligned SDFs averaged across all possible stimulus durations.

#### Neuron selection

In order to be considered for analyzes, individual DLS neurons were required to have displayed an average firing rate during the presentation of the second stimulus of at least 1 Hz (554/602 neurons), and to have been recorded for at least one trial of all possible combinations of T2 and I1 (567/602 neurons), and T2 and I2 (588/602 neurons). An additional set of 42 neurons were deemed unstable by visual inspection of spike rasters and SDFs aligned to several task events. Altogether, these criteria resulted in a selection of 485 out of the original 602 neurons recorded in the delayed duration comparison task variant. Note that unlike the authors, we used the same selection criteria for all reported analyzes.

#### Modulation assessment

In order to quantify the proportion of neurons that were modulated by several behavioral (e.g., choice on the current and previous trials) and experimental factors (e.g., I1, T1, I2 and T2 on the current trial) across the different task epochs, we used these as regressors in multivariable linear regression to fit each neuron’s spike counts recorded in a 334-ms time window aligned to multiple task events (e.g., trial initiation, onset and offset of S1 and S2). For every neuron and every task epoch, the p-values for all factor coefficients were corrected for multiple comparisons using the Benjamini-Hochberg procedure. Lastly, we quantified the proportion of significant coefficients at two significance levels (*α* = 0.01 and *α* = 0.05) for each factor across epochs.

#### Temporal tuning

The temporal tuning of each neuron was estimated separately for each epoch in which the authors enforced their criteria for classifying neurons as ramping or non-ramping(1-s windows centered on the onset and offset of S1 and S2), and stimulus epoch (1-s windows aligned to the onset of S1 and S2), as the center of mass of its baseline-subtracted cross-trial average spike density function.

#### Stereotypy coefficient

For each neuron and within each stimulus epoch, we calculated the Pearson correlation coefficient between two cross-trial average spike density functions, each computed using randomly selected but complementary halves of all its single-trial responses recorded during stimulus presentation. We repeated this procedure 100 times and took the average correlation over permutations to be the stereotypy coefficient for that neuron and stimulus.

#### Firing rate range

A stimulus-specific firing rate range was defined as the difference between the maximum and minimum of each neuron’s cross-trial average spike density function during that stimulus.

### Population analyzes

#### Low dimensional representations of population state

We used PCA to enable visualization of stimulus-aligned population responses in a representative subspace conditioned on task variables of interest. Briefly, we first averaged stimulus-aligned SDFs across all trials for a given neuron and concatenated all the resulting average responses into an *N* × *T* matrix, where *N* is the total number of neurons and *T* is the number of 1-ms time bins. After normalizing these reference data to have zero mean and unit standard deviation along the temporal dimension, we used PCA to find the three orthogonal directions that captured the most variance in stimulus-aligned DLS dynamics. Lastly, when considering a given contrast (e.g., I2), each condition’s trial-averaged population activity was first z-scored using the means and standard deviations estimated from the aforementioned condition-agnostic reference data and then projected onto each of the first three PCs separately, or onto a 2D subspace consisting of an informative section through the space defined by the first three PCs.

#### Neurometric curves

We used linear discriminant analysis (LDA) to find the hyperplane that best separated population state vectors recorded just before the offset of the second stimulus depending on whether T2 was larger or smaller than T1. Briefly, in order to combine trial-by-trial responses of units that were largely recorded separately, rather than simultaneously, pseudo-trial population state vectors were first constructed by concatenating random samples (drawn uniformly and without replacement) from each neuron’s distribution of spike counts in the last 112 ms preceding stimulus offset. This was done separately (but with cross-validation) for each training and test conditions. The former consisted of all possible combinations of T1 and T2 wherein both I1 and I2 were equal to 64 mm s^−1^ (middle intensity), and the latter varied depending on which contrast was used: for I2, test conditions were defined as all possible combinations of T1, T2 and I2 for which I1 = 64 mm s^−1^; for I1, combinations of T1, T2, and I1 for which I2 = 64 mm s^−1^ were used; and for choice, test pseudo-trials were combinations of T1 and T2 for which I1 = I2 = 64 mm s^−1^. We then used LDA to train a classifier to distinguish training pseudo-trials according to what choice they corresponded to, and then used it to “judge” test pseudo-trials. Lastly, for each contrast condition, we fit these “neural judgments” with a 4-parameter psychometric function, analogously to what we did for behavioral ones (see above).

#### Time decoding

We used a naïve (flat prior) Bayes decoder to compute probability distributions over elapsed time using a repeated 2-fold cross-validation scheme with condition-averaged population responses over 100 repetitions.

Briefly, for each repetition we:

1. Discretized time *t* into *B* 1-ms bins, such that *b* = 1, …, *B* and *t*_*b*_ ∈ [− 240, 481] ms relative to the onset of the stimulus of interest;
2. Computed each neuron’s average spike count *n*_*i*_ through time *t* across trials for the training condition and all test conditions using 2-fold cross-validation:

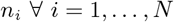
3. Computed each neuron’s tuning curve as a function of time *r*_*i*_(*t*) by convolving its average spike counts through time recorded during the training condition trials with a Gaussian kernel *g*(*t, μ, σ*) with mean *μ* = 0 and standard deviation *σ* = 50 ms:

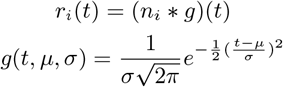
4. Assumed a Poisson spike generation model such that:

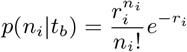
5. Made conditional independence assumptions about neurons, regardless of whether or not they were recorded simultaneously:

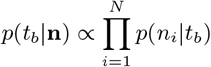

Where **n** = (*n*_1_, *n*_2_, …, *n*_*N*_) is a to-be-decoded test instance of concatenated population activity recorded at a known condition and point in time since the onset of the stimulus of interest;
6. Used Bayes’ rule to create a decoding model that linearly combined all individual neuron encoding models with a uniform prior over decoded time *p*(*t*_*b*_) = 1*/B* ∀ *b* = 1, …, *B*:

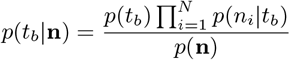

Where the probability *p*(**n**) for the population vector **n** to occur does not have to be estimated explicitly, as it indirectly follows from normalizing the posterior distribution *p*(*t*_*b*_ | **n**) such that it sums to 1 over all possible decoder outcomes, i.e., elapsed time as decoded from DLS ongoing activity;
7. Computed point estimates of decoded time (*μ*_*b*_), accuracy (⟨*ε*⟩) and precision (⟨*σ*⟩):

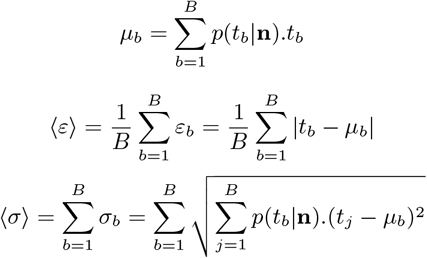

### Simulations

#### Binary choices

To simulate agents driven by nominal NTD and NID, we sampled choices on a trial-by-trial basis from Bernoulli processes according to:

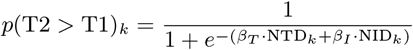

Where *p*(T2 *>* T1)_*k*_, NTD_*k*_ and NID_*k*_ are the Bernoulli process parameter, normalized time difference and normalized intensity difference on trial *k*, respectively. The coefficients *β*_*T*_ and *β*_*I*_ were chosen such that the overall proportion of simulated choices as a function of, and grouped by NTD and NID (Fig. S2B, E left) were similar to those observed in rats (Fig. 1D, G left).

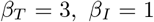

Lastly, to ensure that stimulus and trial statistics were matched between simulated and real data, the trial-wise T1, T2, I1 and I2 values (that determined NTD_*k*_ and NID_*k*_) used in simulations were dictated by the real data.

#### Scalar time estimates

To illustrate how averaging decoded time estimates conditioned on the outcome of choices would mask underlying systematic biases in timekeeping, we simulated marginal distributions over internal time estimates for each stimulus duration assuming scalar timing with a Weber fraction *w* = 0.2. Specifically, internal time distributions were computed by convolving Dirac delta distributions shifted by each possible stimulus duration with Gaussian kernels of increasing standard deviation (*σ* = *w*× *t*). Lastly, when computing joint distributions of T1 and T2 estimates across stimulus pairs, these were assumed to be independent.

#### Spikes times

In three separate instances, we resorted to analyzing spike data simulated from inhomogeneous Poisson point processes on a trial by trial basis (i.e., with a time-varying generative spike rate *λ*_*n,k*_(*t*) for neuron *n* on trial *k*), in a manner that matched (either roughly or exactly depending on the instance) the statistics of real data collected by Toso et al. in terms of numbers of neurons, numbers and types of trials per neuron, and firing statistics. SDFs were estimated from simulated spike trains exactly as described for real data in the spike density functions subsection of the single neuron analyzes section above.

When estimating false positive and false negative rates of our regression-based method for identifying I2-selective neurons during S2 presentation, we generated two sets of surrogate data: a positive and a negative control, with 100% and 0% of neurons modulated by I2, respectively. In the negative control case, for each simulated neuron and on each simulated trial, *λ*_*n,k*_(*t*) was set to the (real) mean S2-aligned SDF of that neuron averaged over middle-I2 trials (I2 = 64 mm s^−1^). For the positive control, the same was true except that generative rates were multiplied by a gain factor proportional to each trial’s I2 (*λ*_*k*_(*t*) = ln (I2_*k*_)*/* ln (64)) before sampling spike times.

When demonstrating that the authors’ operational definitions of ramping and non-ramping neurons predetermined them finding that time was better decoded from the activity of ramps than non-ramps, we simulated populations of Gaussian bumps with three independent parameters: a temporal tuning parameter controlling the average peak time of the simulated bump (*μ*); a stereotypy parameter controlling trial-to-trial variability in that peak time (*η*); and a gain parameter controlling peak amplitude relative to baseline (*γ*).

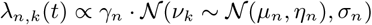

To assess if variability in each parameter on its own was sufficient to drive performance differences in time decoding between ramps and non-ramps, we simulated four different populations of our model bumps that differed with respect to which parameters were held constant and which were sampled from arbitrary distributions (informed by the biases observed in real data):

1. *μ* ∼ 𝒰 [−500, 1500] ms, *η* = 0 ms, *γ* = 3 s^−1^
2. *μ* ∼ 𝒰 [−500, 1500] ms, *η* ∼ Exp(250) ms, *γ* = 3 s^−1^
3. *μ* ∼ 𝒰 [−500, 1500] ms, *η* = 0 ms, *γ* ∼ Exp(3) s^−1^
4. *μ* ∼ 𝒰 [−500, 1500] ms, *η* ∼ Exp(250) ms, *γ* ∼ Exp(3) s^−1^

Lastly, when generating ground truth data to validate our clusterability analysis, we adopted a hybrid approach wherein the subset of DLS neurons that had been labeled as ramps using the authors’ classification criteria were replaced with simulated ramps proper, equally split among up- and down-ramping ones. Importantly, simulated ramps inherited the trial (number and type) and firing (baseline and range) statistics of the real neurons they replaced.

### Statistics

Departures from unimodality were assessed using Hartigan’s dip test (Hartigan & Hartigan, 1985), and differences between distributions were assessed using two-sample Kolmogorov-Smirnov tests with Bonferroni correction for multiple comparisons (where applicable).

In all cases, test outcomes were denoted as: NS, *p* ≥ 0.05; *, *p <* 0.05; **, *p <* 0.01, written in the same color as that of the corresponding distribution for unimodality tests, and in black near the horizontal solid black lines plotted above the underlying distributions for two-sample tests.

## Supplementary figures

**Figure S1:**
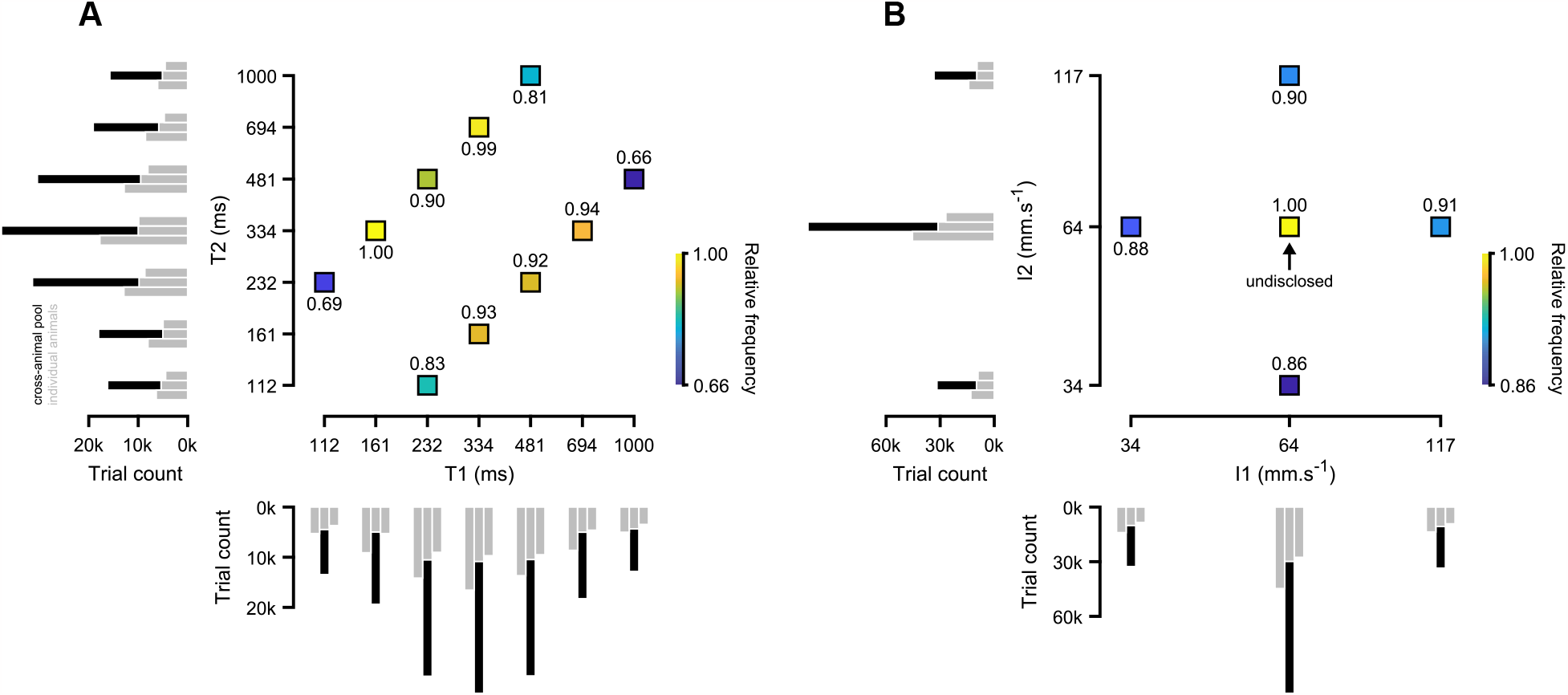
Distribution of stimulus features in the delayed duration comparison task. (**A**) Main axes: frequency of stimulus duration pairs (T1 and T2) relative to the most common pair (T1 = 161; T2 = 334 ms). Left: Distribution of sampled T2 durations for individual DLS-implanted animals (in gray) and pooling across them (black). (**B**) Same as (A), but for stimulus intensity. The black arrow in the main axes highlights the intensity pair that was undisclosed in both the original study by Toso, Reinartz, et al., 2021 and its erratum (I1 = 64; I2 = 64 mm s^−1^).

**Figure S2:**
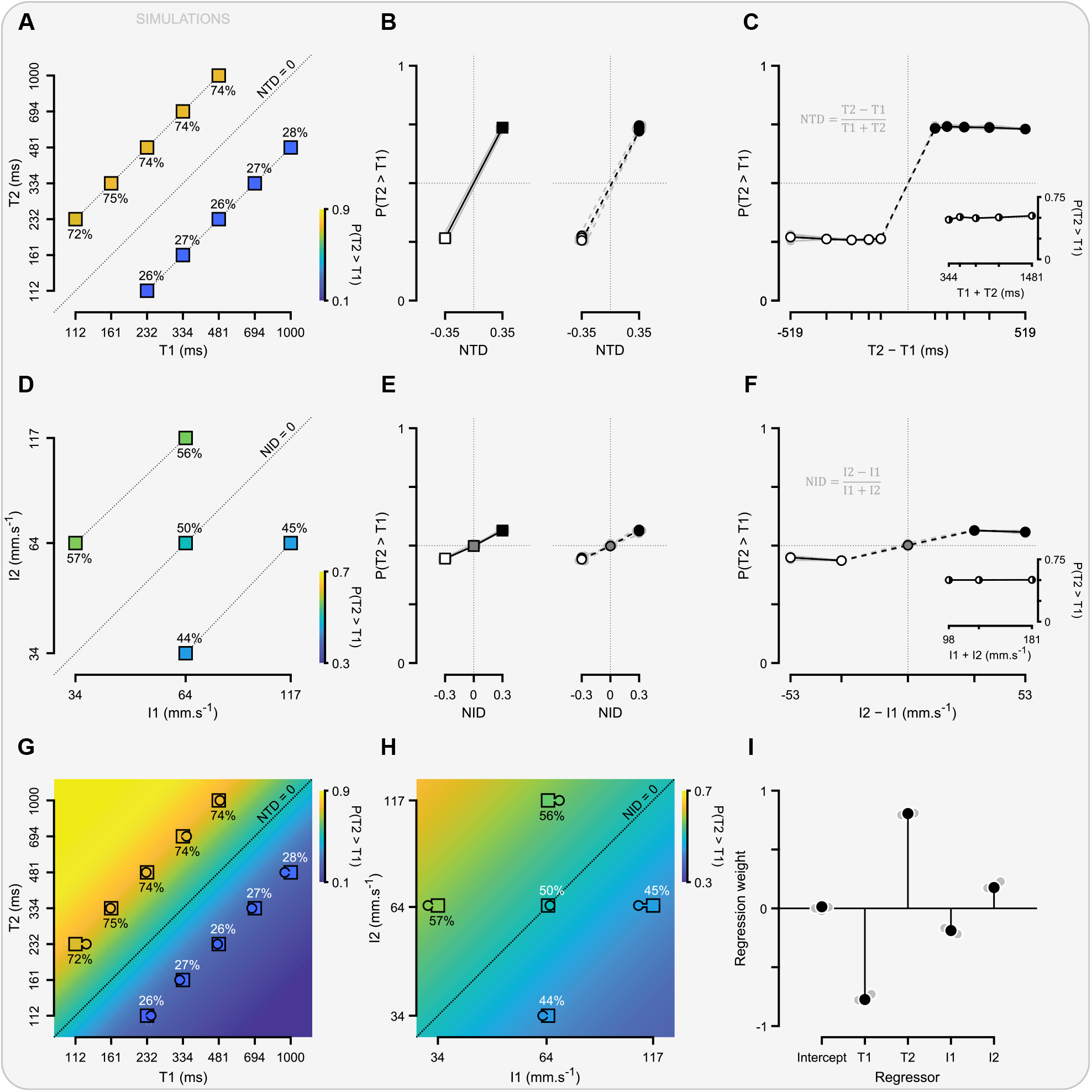
Performance of simulated agents strictly driven by NTD and biased by NID. (**A**) Proportion of T2 > T1 choices for all stimulus duration pairs pooled across three simulated agents. Dotted lines above, below and at the identity line represent NTD levels of +0.35, −0.35 and 0, respectively. (**B**) Proportion of T2 > T1 choices as a function of NTD and grouped by NTD level (left) or stimulus pair (right). Gray markers and lines refer to individual agents, and black and white ones to cross-agent averages for each NTD level. Dashed lines on the right connect the two stimulus pairs with smaller absolute duration differences (|T2 − T1|). (**C**) Proportion of T2 > T1 choices conditioned on all possible values of the numerator (main axes) and denominator (inset) terms in the definition of NTD. (**D-F**) Same as (A-C), but for stimulus intensity and NID. (**G**) Same data as in (A), but overlaying the performance rule that generated them. For each duration pair, a line was drawn between its T1 and T2 coordinates and the horizontally shifted point for which generative performance was closest to simulated performance on that pair. (**H**) Similar to (G), but for stimulus intensity. (**I**) Coefficients of a logistic regression model fit to simulated choice data using all four stimulus features (T1, T2, I1 and I2) as regressors. Gray markers refer to fits for individual agents, and black ones to a fit pooling data across all three agents.

**Figure S3:**
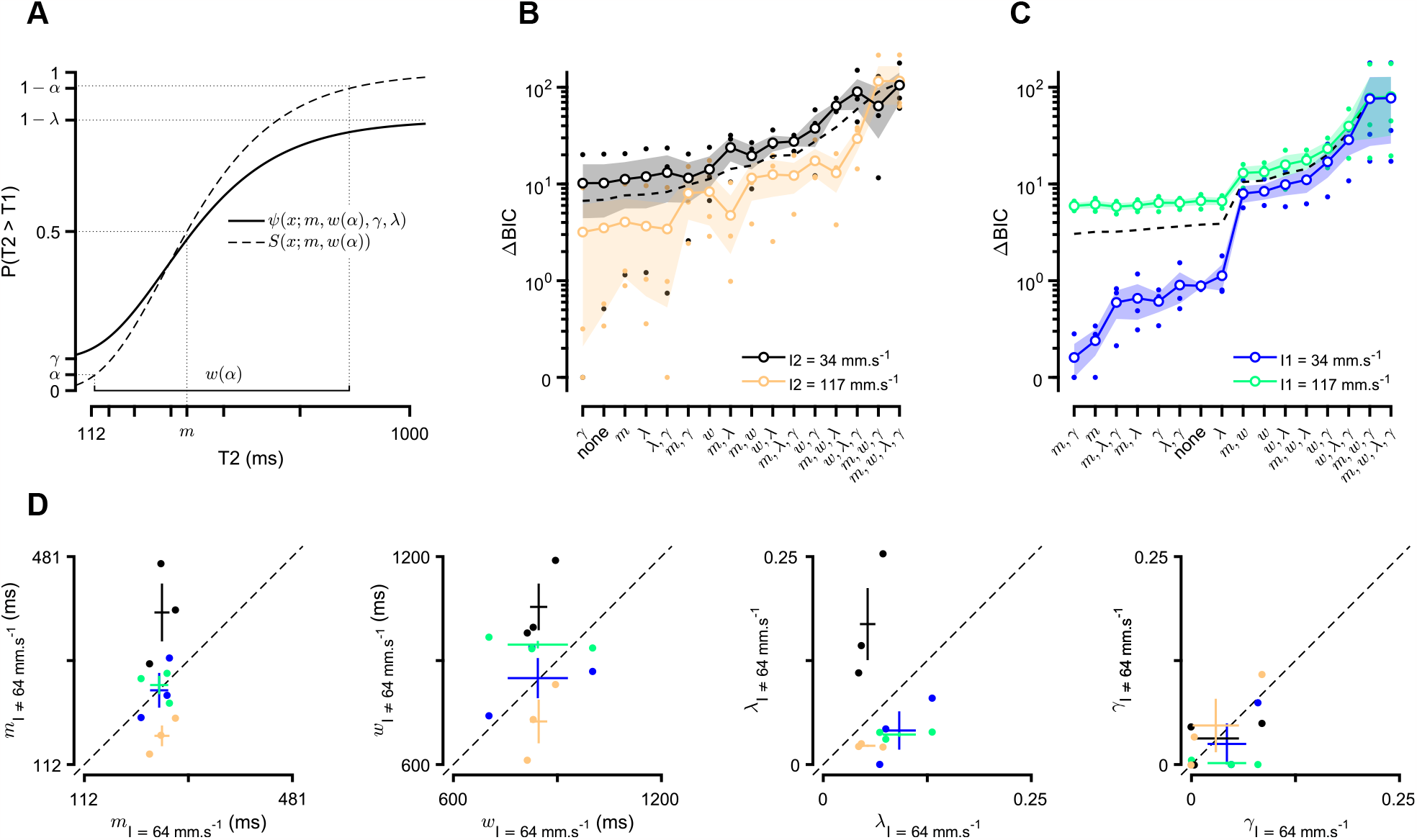
Multiple parameters of the psychometric function were affected by the intensity of the second stimulus, but not that of the first. (**A**) Schematic illustrating the parameterization of the psychometric function used throughout (adapted from Schütt et al., 2016). (**B**) Model comparison between psychometric fits to choices of individual rats in middle-I2 (I2 = 64 mm s^−1^) and extreme-I2 trials (high: I2 = 117 mm s^−1^; low: I2 = 34 mm s^−1^), measured as the difference between each model’s BIC score and that of the rat-specific model with the lowest BIC across low and high intensities. Extreme-I2 models are labeled according to which parameters of the psychometric function were free and which were set to those of the corresponding middle-I2 fits (e.g., “none” means that all parameters were free, and “*m*” means that only the threshold parameter was fixed). Small and large markers correspond to individual animals and cross-animal averages (mean ± SEM), respectively. The dashed black line corresponds to the I2-agnostic cross-animal average, according to which models were sorted along the *x*-axis. (**C**) Same as (B), but for I1. (**D**) Point estimates for each of the four parameters of the psychometric function introduced in (A) estimated using only middle-intensity (*x*-axis) or extreme-intensity trials (*y*-axis). Small circles correspond to individual animals, and crosses to cross-animal averages (mean ± SEM).

**Figure S4:**
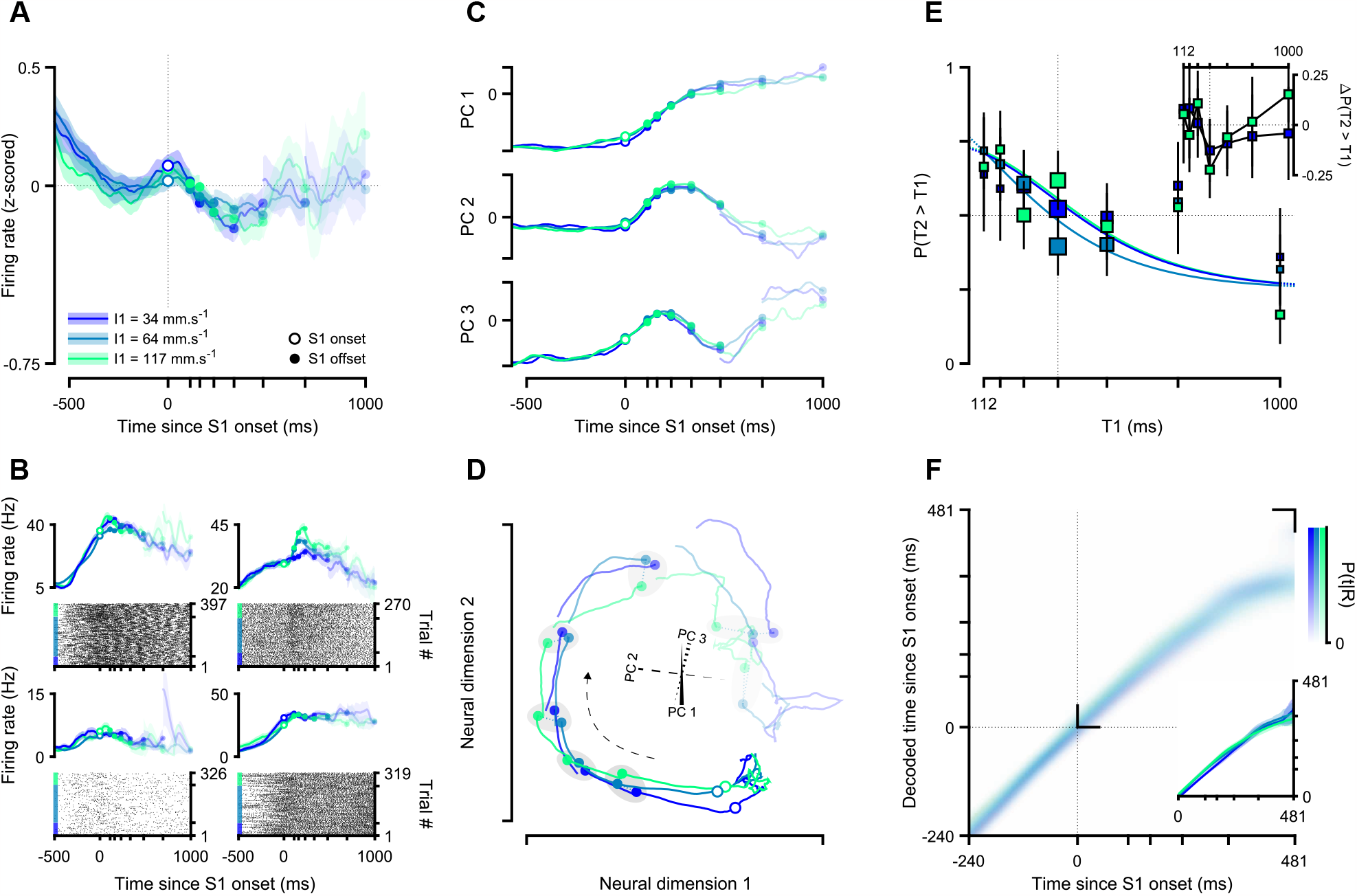
DLS dynamics recorded during the first stimulus were negligibly affected by the intensity of the first stimulus. (**A**) Average normalized population activity conditioned on I1 and aligned to the onset of S1 (mean ± SEM across neurons). (**B**) Responses of four example DLS neurons during S1. For each neuron, spike density functions (mean ± SEM across trials) are conditioned on I1 and plotted above its spike raster, where trials are first sorted by I1, and then by T1 within each I1 level. (**C**) I1-split population activity projected onto each of the first three PCs computed using I1-agnostic population activity aligned to the onset of S1. (**D**) Population activity conditioned on I1 projected onto a 2D section of the subspace defined by the three PCs shown in (C). The 3-axis gizmo at the center depicts the projections of the axes defined by the first three PCs onto the chosen 2D section, the dashed black arrow indicates the direction of time, and the gray patches group population states recorded at each possible stimulus offset across I1. (**E**) Neurometric curves trained using S1-aligned population activity from middle-I1 trials conditioned on I1 (main axes), and differences between the proportion of T2 > T1 neural “reports” in high (I1 = 117 mm s^−1^) and low (I1 = 34 mm s^−1^) intensity trials compared to middle intensity (I1 = 64 mm s^−1^) trials (inset, mean ± propagated SEM). (**F**) Time estimates decoded from DLS activity around S1 presentation conditioned on I1. Main axes: mean posterior distributions across runs. Inset: MAP point estimates averaged across runs (median and IQR). The opacity of lines, markers and patches in (A-D) reflects the relative number of trials contributing to the underlying estimates.

**Figure S5:**
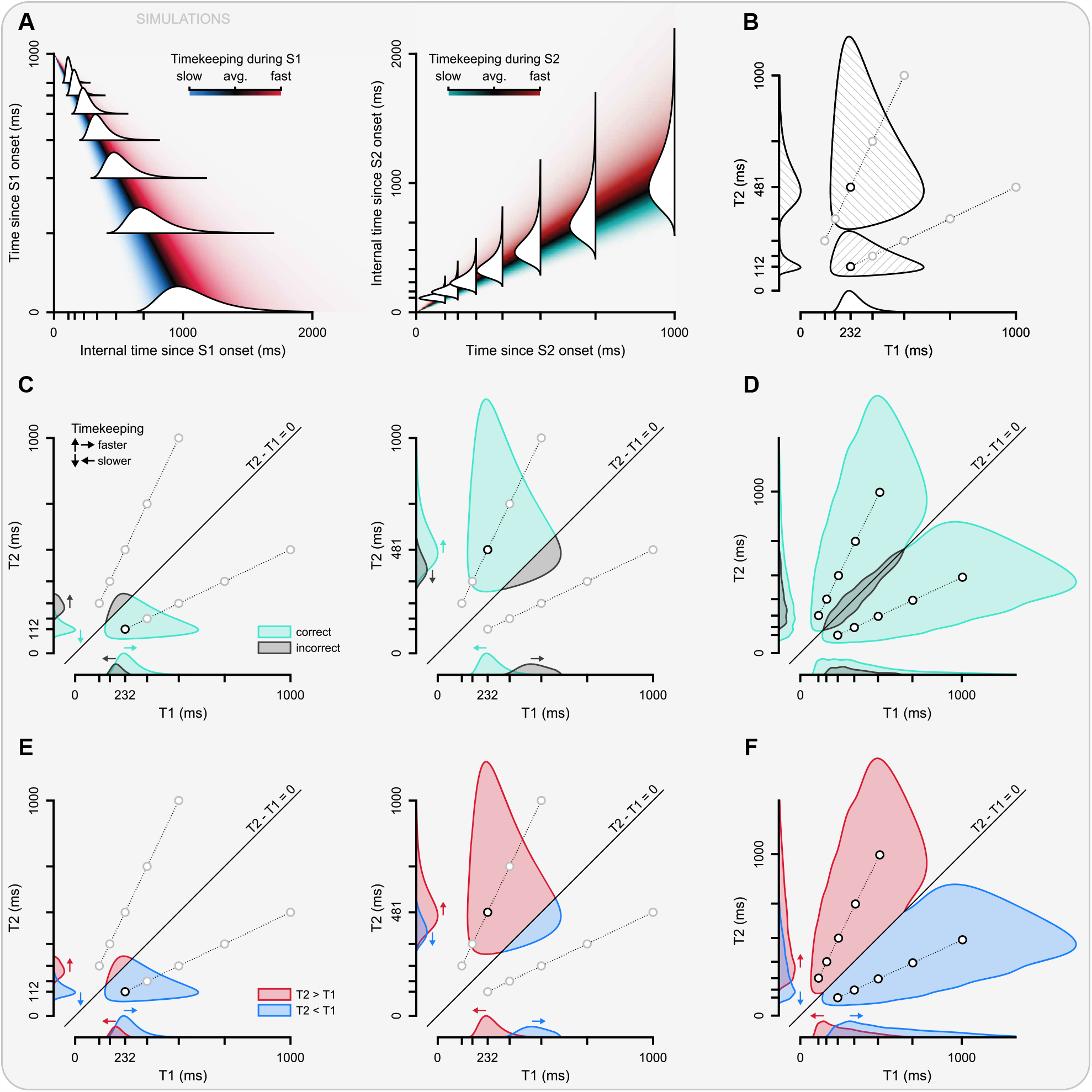
Systematic biases in timekeeping across stimulus pairs are obscured (revealed) by conditioning trials on the outcome (category) of choices. (**A**) Hypothetical distributions of internal estimates of elapsed time as a function of experimental time since the onset of the first (left) and second (right) stimulus, plotted both continuously (colored gradient in the background) and discretely at the set of possible stimulus offset times (white distributions in the foreground). (**B**) Joint distributions of internal estimates of T1 and T2 for two pairs of stimuli with their respective marginals projected on each axis. (**C**) Joint and marginal distributions of T1 and T2 estimates for a stimulus pair for which T2 was shorter (left) or larger (right) than T1, conditioned on whether a direct comparison of those estimates would produce correct or incorrect choices. (**D**) Same as (C) but for all stimulus pairs. (**E-F**) Same as (C-D) but conditioned on the category of hypothetical choices strictly informed by a comparison of T1 and T2 estimates.

## Supplementary text

Here we show that rats’ performance in the delayed duration comparison task was not driven by the nominal NTD and biased by NID as idealized and reported by Toso and colleagues. Instead, we find a marked asymmetry in the contributions of the two stimuli onto choices, with the second vibration’s duration dominating the comparison. Moreover, vibration intensity of the first stimulus was virtually inconsequential to animals’ decisions, meaning that the reported intensity bias was driven almost exclusively by the intensity of the second stimulus. All else being equal, failure to account for this might have misled the authors into giving equal emphasis to both stimuli when analyzing and interpreting DLS activity, which on its own would not support the central claims of Toso et al. 2021. However, this inaccuracy was aggravated by a pervasive preprocessing error that resulted in all spike times being shifted 200 ms into the future relative to behavioral events. Though this constitutes a substantial misalignment error, especially considering the range of durations used for stimulus ([112, 1000] ms) and other delay epochs ([500, 2000] ms) in the task, it again would not, on its own, have accounted for the central claims of Toso and colleagues. Those can only be fully understood when combined with arbitrary and convoluted operational definitions (e.g., those used to identify ramping and intensity-modulated neurons), ill-posed analyzes regarding the use of nominal choice outcome (i.e., correctness) as a contrast (detailed in the main text), and a cross-epoch contamination issue affecting several analyzes of stimulus-aligned neural data. Specifically, in multiple analyzes aimed at characterizing DLS activity during stimulus presentation (e.g., when assessing intensity modulation at the single-neuron and population levels), a window of 1000 ms aligned on stimulus (first or second) onset was used irrespective of the trial-specific durations of the stimuli in question, meaning that for stimulus durations shorter than 1000 ms, activity from the subsequent epoch(s) (likely including periods in which the animals were already moving for shorter stimuli) contaminated the analysis.

We must note that Toso et al. have since published an erratum in which they partially and selectively addressed two of these issues, despite us sharing earlier versions of all of the analyzes (figures and implementation code) presented in this manuscript. Specifically, the authors focused their correction on the issues of spike misalignment and cross-epoch contamination. The former, though striking, was on its own of little consequence in the authors’ interpretation of the data. The latter, though a much bigger determinant of the authors’ results and central conclusions, was deemphasized as a potential source of confounds and framed as a legitimate design choice in light of constraints imposed by data scarcity. More importantly though, in their revised summary the authors maintained all of their original claims, only complemented by an addendum acknowledging the modulation of DLS activity by the intensity of the second but not that of the first stimulus, but without an accompanying acknowledgement of the asymmetric contribution of these stimulus features onto animals’ behavior.

For these reasons, we consider the existing correction to be insufficient and ineffectual, given that it exists only as a separate publication that pales in visibility in comparison to the original, undoctored publication.

## Acronyms

BIC: Bayesian information criterion. 19, 27
DLS: dorsolateral striatum. 1–12, 14, 15, 17, 19–22, 25, 28, 30
GLM: generalized linear model. 2
I1: intensity of the first stimulus. 2–4, 6–9, 11, 12, 14, 19–21, 25–28
I2: intensity of the second stimulus. 2–8, 11, 12, 14, 19–22, 25–27
IQR: interquartile range. 7, 9, 10, 14, 17, 28
ISI: inter stimulus interval. 2, 8, 11, 12
LDA: linear discriminant analysis. 20
MAP: *maximum a posteriori*. 7, 9, 10, 28
MSN: medium spiny neuron. 11
NID: normalized intensity difference. 3, 4, 21, 26, 30
NTD: normalized time difference. 3, 4, 6, 21, 26, 30
PC: principal component. 5, 7–10, 15, 17, 20, 28
PCA: principal component analysis. 5, 8, 20
S1: first stimulus. 2, 3, 8, 11, 12, 14, 17, 19, 20, 28
S2: second stimulus. 2, 5, 7–12, 14, 15, 17, 19, 20, 22
SD: standard deviation. 17
SDF: spike density function. 19, 20, 22
SEM: standard error of the mean. 6, 7, 9, 10, 27, 28
T1: duration of the first stimulus. 2–9, 11, 12, 19–22, 25, 26, 28, 29
T2: duration of the second stimulus. 2–12, 19–22, 25, 26, 28, 29
V1: primary visual cortex. 2

## References

Adolfsson, A., Ackerman, M., & Brownstein, N. C. (2019). To cluster, or not to cluster: An analysis of clusterability methods. Pattern Recognit., 88, 13–26.

Akrami, A., Kopec, C. D., Diamond, M. E., & Brody, C. D. (2018). Posterior parietal cortex represents sensory history and mediates its effects on behaviour. Nature, 554 (7692), 368–372.

Ashourian, P., & Loewenstein, Y. (2011). Bayesian inference underlies the contraction bias in delayed comparison tasks.

Bakhurin, K. I., Goudar, V., Shobe, J. L., Claar, L. D., Buonomano, D. V., & Masmanidis, S. C. (2017). Differential encoding of time by prefrontal and striatal network dynamics. J. Neurosci., 37 (4), 854–870.

Boboeva, V., Pezzotta, A., Clopath, C., & Akrami, A. (2023, January). From recency to central tendency biases in working memory: A unifying network model.

Coull, J. T., Vidal, F., Nazarian, B., & Macar, F. (2004). Functional anatomy of the attentional modulation of time estimation. Science, 303 (5663), 1506–1508.

Cruz, B. F., Guiomar, G., Soares, S., Motiwala, A., Machens, C. K., & Paton, J. J. (2022). Action suppression reveals opponent parallel control via striatal circuits. Nature, 607 (7919), 521–526.

Donnelly, N. A., Paulsen, O., Robbins, T. W., & Dalley, J. W. (2015). Ramping single unit activity in the medial prefrontal cortex and ventral striatum reflects the onset of waiting but not imminent impulsive actions. Eur. J. Neurosci., 41 (12), 1524–1537.

Emmons, E. B., De Corte, B. J., Kim, Y., Parker, K. L., Matell, M. S., & Narayanan, N. S. (2017). Rodent medial frontal control of temporal processing in the dorsomedial striatum. J. Neurosci., 37 (36), 8718–8733.

Fassihi, A., Akrami, A., Esmaeili, V., & Diamond, M. E. (2014). Tactile perception and working memory in rats and humans. Proc. Natl. Acad. Sci. U. S. A., 111 (6), 2331–2336.

Gibbon, J. (1991). Origins of scalar timing. Learn. Motiv., 22 (1), 3–38.

Gouvêa, T. S., Monteiro, T., Motiwala, A., Soares, S., Machens, C., & Paton, J. J. (2015). Striatal dynamics explain duration judgments. Elife, 4.

Hartigan, J. A., & Hartigan, P. M. (1985). The Dip Test of Unimodality. The Annals of Statistics, 13 (1), 70–84. 10.1214/aos/1176346577

Hernández, A., Salinas, E., García, R., & Romo, R. (1997). Discrimination in the sense of flutter: New psychophysical measurements in monkeys. J. Neurosci., 17 (16), 6391–6400.

Hollingworth, H. L. (1910). The central tendency of judgment. The Journal of Philosophy, Psychology and Scientific Methods, 7 (17), 461–469.

Jazayeri, M., & Shadlen, M. N. (2010). Temporal context calibrates interval timing. Nat. Neurosci., 13 (8), 1020–1026.

Jin, D. Z., Fujii, N., & Graybiel, A. M. (2009). Neural representation of time in cortico-basal ganglia circuits. Proc. Natl. Acad. Sci. U. S. A., 106 (45), 19156–19161.

Kim, J., Kim, D., & Jung, M. W. (2018). Distinct dynamics of striatal and prefrontal neural activity during temporal discrimination. Front. Integr. Neurosci., 12, 34.

Kobak, D., Brendel, W., Constantinidis, C., Feierstein, C. E., Kepecs, A., Mainen, Z. F., Qi, X.-L., Romo, R., Uchida, N., & Machens, C. K. (2016). Demixed principal component analysis of neural population data. Elife, 5.

Lau, B., & Glimcher, P. W. (2008). Value representations in the primate striatum during matching behavior. Neuron, 58 (3), 451–463.

Malapani, C., Rakitin, B., Levy, R., Meck, W. H., Deweer, B., Dubois, B., & Gibbon, J. (1998). Coupled temporal memories in parkinson’ s disease: A dopamine-related dysfunction. J. Cogn. Neurosci., 10 (3), 316–331.

Malapani, C., & Fairhurst, S. (2002). Scalar timing in animals and humans. Learn. Motiv., 33 (1), 156–176.

Mante, V., Sussillo, D., Shenoy, K. V., & Newsome, W. T. (2013). Context-dependent computation by recurrent dynamics in prefrontal cortex. Nature, 503 (7474), 78–84.

Maricq, A. V., & Church, R. M. (1983). The differential effects of haloperidol and methamphetamine on time estimation in the rat. Psychopharmacology, 79 (1), 10–15.

Matell, M. S., Meck, W. H., & Nicolelis, M. A. L. (2003). Interval timing and the encoding of signal duration by ensembles of cortical and striatal neurons. Behav. Neurosci., 117 (4), 760–773.

Meck, W. H. (2006). Neuroanatomical localization of an internal clock: A functional link between mesolimbic, nigrostriatal, and mesocortical dopaminergic systems. Brain Res., 1109 (1), 93–107.

Mello, G. B. M., Soares, S., & Paton, J. J. (2015). A scalable population code for time in the striatum. Curr. Biol., 25 (9), 1113–1122.

Monteiro, T., Rodrigues, F. S., Pexirra, M., Cruz, B. F., Gonçalves, A. I., Rueda-Orozco, P. E., & Paton, J. J. (2023). Using temperature to analyze the neural basis of a time-based decision. Nat. Neurosci.

Paton, J. J., & Buonomano, D. V. (2018). The neural basis of timing: Distributed mechanisms for diverse functions. Neuron, 98 (4), 687–705.

Rao, S. M., Mayer, A. R., & Harrington, D. L. (2001). The evolution of brain activation during temporal processing. Nat. Neurosci., 4 (3), 317–323.

Reynolds, J. N., Hyland, B. I., & Wickens, J. R. (2001). A cellular mechanism of reward-related learning. Nature, 413 (6851), 67–70.

Rigotti, M., Barak, O., Warden, M. R., Wang, X.-J., Daw, N. D., Miller, E. K., & Fusi, S. (2013). The importance of mixed selectivity in complex cognitive tasks. Nature, 497 (7451), 585–590.

Romo, R., Brody, C. D., Hernández, A., & Lemus, L. (1999). Neuronal correlates of parametric working memory in the prefrontal cortex. Nature, 399 (6735), 470–473.

Rowe, K. C., Paulsen, J. S., Langbehn, D. R., Duff, K., Beglinger, L. J., Wang, C., O’Rourke, J. J. F., Stout, J. C., & Moser, D. J. (2010). Self-paced timing detects and tracks change in prodromal huntington disease. Neuropsychology, 24 (4), 435–442.

Samejima, K., Ueda, Y., Doya, K., & Kimura, M. (2005). Representation of action-specific reward values in the striatum. Science, 310 (5752), 1337–1340.

Schultz, W., Dayan, P., & Montague, P. R. (1997). A neural substrate of prediction and reward. Science, 275 (5306), 1593–1599.

Schütt, H. H., Harmeling, S., Macke, J. H., & Wichmann, F. A. (2016). Painfree and accurate bayesian estimation of psychometric functions for (potentially) overdispersed data. Vision Res., 122, 105–123.

Soares, S., Atallah, B. V., & Paton, J. J. (2016). Midbrain dopamine neurons control judgment of time. Science, 354 (6317), 1273–1277.

Steiner, H., & Tseng, K. Y. (2016, September). Handbook of basal ganglia structure and function. Academic Press.

Tai, L.-H., Lee, A. M., Benavidez, N., Bonci, A., & Wilbrecht, L. (2012). Transient stimulation of distinct subpopulations of striatal neurons mimics changes in action value. Nat. Neurosci., 15 (9), 1281–1289.

Toso, A., Fassihi, A., Paz, L., Pulecchi, F., & Diamond, M. E. (2021). A sensory integration account for time perception. PLoS Comput. Biol., 17 (1), e1008668.

Toso, A., Reinartz, S., Pulecchi, F., & Diamond, M. E. (2021). Time coding in rat dorsolateral striatum. Neuron, 109 (22), 3663–3673.e6.

Treisman, M. (1963). Temporal discrimination and the indifference interval: Implications for a model of the “internal clock”. Psychological Monographs: General and Applied, 77 (13), 1–31.

Urai, A. E., Braun, A., & Donner, T. H. (2017). Pupil-linked arousal is driven by decision uncertainty and alters serial choice bias. Nat. Commun., 8, 14637.

Wang, J., Narain, D., Hosseini, E. A., & Jazayeri, M. (2017). Flexible timing by temporal scaling of cortical responses. Nat. Neurosci.

Wolff, S. B. E., Ko, R., & Ölveczky, B. P. (2022). Distinct roles for motor cortical and thalamic inputs to striatum during motor skill learning and execution. Sci Adv, 8 (8), eabk0231.

Xiang, Y., Graeber, T., Enke, B., & Gershman, S. J. (2021). Confidence and central tendency in perceptual judgment. Atten. Percept. Psychophys., 83 (7), 3024–3034.

